# Human brain region-specific variably methylated regions (VMRs) are enriched for heritability of distinct neuropsychiatric traits

**DOI:** 10.1101/2021.01.02.425010

**Authors:** Lindsay F. Rizzardi, Peter F. Hickey, Adrian Idrizi, Rakel Tryggvadóttir, Colin M. Callahan, Kimberly E. Stephens, Sean D. Taverna, Hao Zhang, Sinan Ramazanoglu, GTEx Consortium, Kasper D. Hansen, Andrew P. Feinberg

## Abstract

**BACKGROUND:** DNA methylation dynamics in the brain are associated with normal development and neuropsychiatric disease and differ across functionally distinct brain regions. Previous studies of genome-wide methylation differences among human brain regions focused on limited numbers of individuals and one to two brain regions.

**RESULTS:** Using GTEx samples, we have generated a resource of DNA methylation in purified neuronal nuclei from 8 brain regions as well as lung and thyroid tissues from 12-23 donors. We identified differentially methylated regions between brain regions (DMRs) among neuronal nuclei in both CpG (181,146) and non-CpG (264,868) contexts, few of which were unique to a single pair-wise comparison. This significantly expands the knowledge of differential methylation across the brain by 10-fold. In addition, we present the first differential methylation analysis among neuronal nuclei from basal ganglia tissues and identified 2,295 unique CpG DMRs, many associated with ion transport. Consistent with prior studies, CpG DMRs were enriched in regulatory regions while non-CpG DMRs were enriched in intergenic regions. We also identified 81,130 regions of variably CpG methylated regions (VMRs), i.e. variable methylation among individuals in the same brain region, which were enriched in regulatory regions and in CpG DMRs. Many VMRs were unique to a specific brain region, with only 202 common across all brain regions, as well as lung and thyroid. VMRs identified in the amygdala, anterior cingulate cortex, and hippocampus were enriched for heritability of schizophrenia.

**CONCLUSIONS:** These data suggest that epigenetic variation in these particular human brain regions could be associated with the risk for this neuropsychiatric disorder.

## INTRODUCTION

DNA methylation patterns are altered throughout development to establish distinct cell fates [1–3]. Analyses of methylation changes during cortical development have shown that, in neurons, these changes are often associated with synaptogenesis during the first 5 years of life [4]. These regions of dynamic methylation have been linked to neuropsychiatric disorders that are thought to have developmental origins [5–8]. In adulthood, the epigenome is intimately involved in neuronal plasticity in response to environmental exposures and synaptic activity [9, 10]. DNA methylation and chromatin features differ among neurons from functionally distinct adult brain regions and these regions are enriched for heritability of neuropsychiatric traits [11, 12]. The PsychENCODE consortium has made substantial contributions to our understanding of gene regulation in the human brain, particularly during development and disease [13, 14], and in distinct cell populations within the cortex [15]. However, these efforts have been somewhat limited either by examination of multiple cell populations within a single cortical region or by the use of bulk tissues that are strongly confounded by cellular heterogeneity. Thus, the extent of DNA methylation variation within neuronal populations from many adult human brain regions remains unknown and is the focus of this work.

The Genotype-Tissue Expression (GTEx) project [16–19] has enabled unprecedented analysis and understanding of tissue-specific expression and the genetic determinants of this expression. To complement the expression data generated by GTEx, the enhanced GTEx project [20] has profiled additional molecular traits across a subset of GTEx samples. We previously explored DNA methylation across four brain regions using non-GTEx samples from six individuals [12]. Here, we describe the results of the enhanced GTEx DNA methylation project that examined 182 samples representing 8 brain regions and 2 somatic tissues from 12-24 GTEx donors with extensive profiling of DNA methylation and the influence of genotype on methylation variability. Moreover, based on our earlier work, purification of neuronal rather than bulk tissue DNA prior to analysis was imperative to obtain high quality methylation data due to the extensive neuron/glia heterogeneity even in adjacent sections from the same brain region. In contrast, GTEx expression data was obtained exclusively from bulk tissue samples confounded by cellular heterogeneity; therefore, we focused on how genotype influenced methylation rather than how methylation impacted gene expression. We identify differentially methylated regions among NeuN+ cells from 8 brain regions in both a CpG and non-CpG context, greatly extending our knowledge of functional epigenetic differences across the brain. Importantly, the larger number of brain regions and individuals enabled our identification of variably methylated regions (VMRs) that we initially defined as regions that are highly variable among individuals within a given tissue type [21, 22]. Surprisingly, many VMRs were brain region-specific, 60-95% involved two or more regions and were concordant for methylation, while only 202 were ubiquitous across all 8 brain regions and 2 somatic tissues. Along with CpG differentially methylated regions (CG-DMRs), VMRs from three brain regions were enriched for heritability of schizophrenia suggesting the importance of epigenetic variation in neuropsychiatric disease risk.

## RESULTS

### Characterizing the neuronal DNA methylation landscape across 8 brain regions

Previous reports from us and others have shown distinct epigenetic landscapes among functionally diverse brain regions [11, 12]. Our previous study was limited to a small number of individuals and brain regions. However, we demonstrated that brain region-specific DNA methylation was primarily present in neuronal rather than non-neuronal nuclei. Further, the ratio of neurons to non-neurons even between adjacent sections from the same brain region differed greatly, severely confounding analysis of differential methylation in non-purified nuclei. Therefore, we applied a strategy of neuronal nuclei purification prior to whole genome bisulfite sequencing. Analyzing a much larger number of individuals and brain regions enabled us to address the potential existence of VMRs (regions of interindividual variation in methylation within a tissue), their relationship to each other, and the relationship to SNPs identified in these GTEx donors. We examined 8 brain regions collected from GTEx donors: amygdala (n = 12), anterior cingulate cortex (BA24) (n = 15), caudate (n = 22), frontal cortex (BA9) (n = 24), hippocampus (n = 20), hypothalamus (n = 13), nucleus accumbens (n = 23), and putamen (n = 16). In addition, we analyzed methylation of DNA isolated from two non-brain tissues from GTEx donors: lung (n = 18) and thyroid (n = 19) for a total of 182 samples (**Supplemental Table S1**). Neuronal nuclei were isolated from brain tissues based on positive NeuN (*RBFOX3*) staining via fluorescence activated nuclei sorting and NeuN+ nuclei are referred to as neuronal, while noting that this fraction is composed of multiple subpopulations (**Fig. S1a**). We generated >30 billion uniquely mapped 150-bp paired-end reads with an average depth >10X post-processing (**Supplemental Table S2)**. Several samples were excluded due to genotype discordance, as the shipped sample genotype did not match the biobank records (**Supplemental Table S3**). Five additional samples were excluded after principal component analysis revealed sample mislabeling prior to receipt by our lab. We confirmed this by determining which tissues most closely matched the methylation of these samples (**Fig. S1b**).

Principal component analysis of global neuronal DNA methylation levels revealed clear segregation of these brain regions in the first two principal components (**Fig. 1a**). We performed a differential analysis of CpG methylation identifying CG-DMRs, i.e. regions of differential CpG methylation among neuronal nuclei isolated from each brain region. Given that a single CG-DMR can represent a difference among multiple brain regions, rather than perform 28 pairwise comparisons, we used an F-test to identify 174,482 statistically significant autosomal neuronal CG-DMRs which are defined as regions of the genome where at least 2 of the 8 brain regions have different levels of CpG methylation (**Supplemental Table S4**). We control the family-wise error rate at 5% by permutation, and we use BSmooth to leverage information from nearby CpGs by smoothing. In a pilot study, we profiled NeuN+ cells from 4 brain regions using whole genome bisulfite sequencing on samples from 6 different individuals not part of GTEx [12]. We find that 99.5% of our previously identified neuronal CG-DMRs (13019/13074) overlap with CG-DMRs from our new analysis of GTEx samples (after correcting for multiple testing). To make a more precise comparison, we examined the correlation between the methylation differences between two tissues as measured separately in Rizzardi et al [12] and this study (tissues were nucleus accumbens and prefrontal cortex, selected because most of the CG-DMRs from Rizzardi et. al. [12]were between those two tissues). We find a striking correlation of 0.97 as shown in Figure S1c highlighting the reproducibility of our experimental and analytical approaches across biobanks. This high level of reproducibility holds even when examining methylation differences among all 13,074 neuronal DMRs identified in [12] between two very similar cortical regions (Figure S1d), which suggests that our approach is conservative, likely because we control the family-wise error rate.

**Figure 1.**
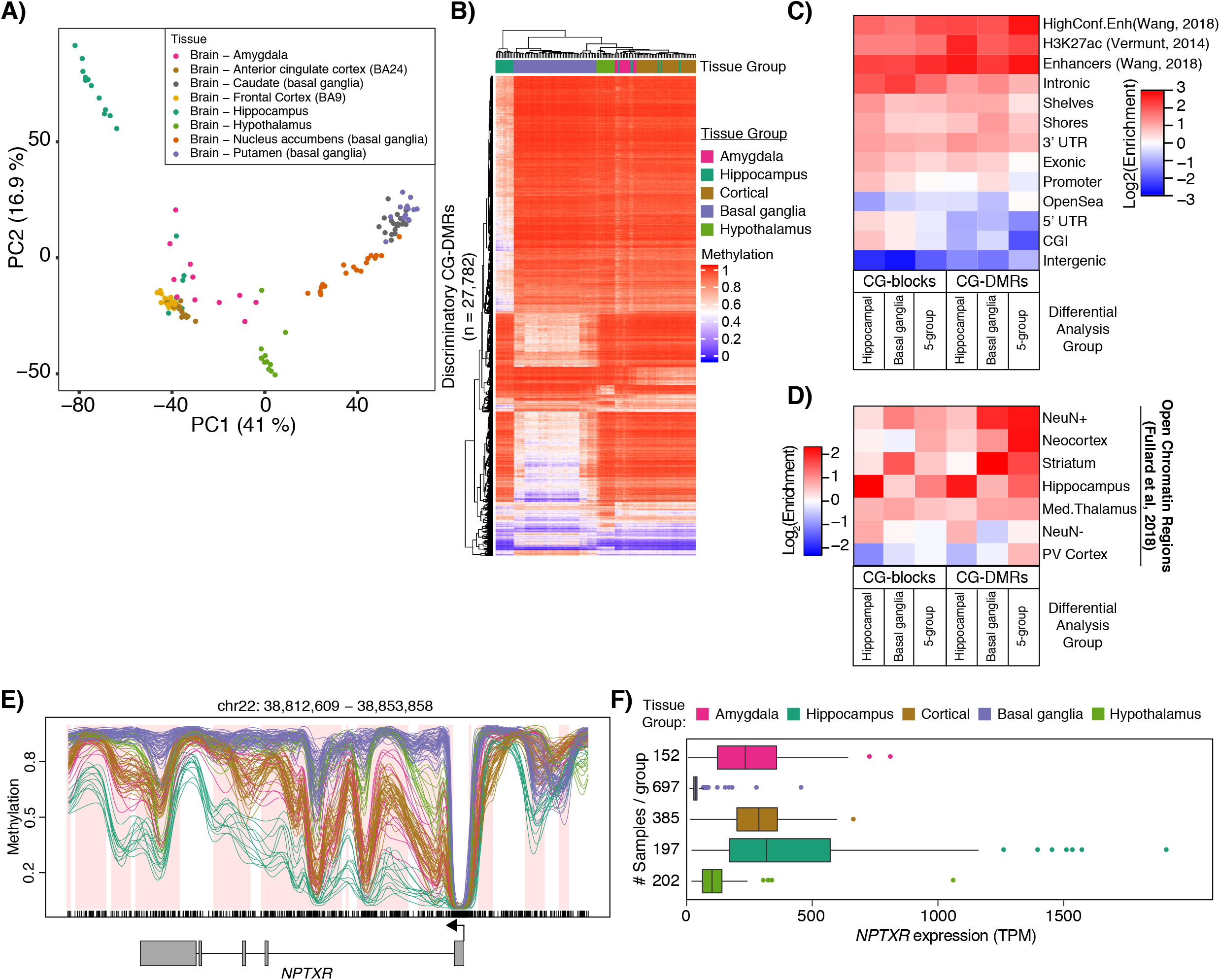
Identification of CG-DMRs among neurons of functionally distinct brain regions. DNA methylation was assessed in neuronal nuclei isolated from 8 brain regions as indicated from 12-24 individuals. A) Principle component analysis of distances derived from average autosomal CpG methylation in 1 kb bins. B) Hierarchical clustering of samples based on the average methylation per sample in the most discriminatory CG-DMRs (see Methods). C) Heatmap representing log2 enrichment of CpGs within CG-DMRs and blocks identified in each CG-DMR analysis compared to the rest of the genome for genomic features. Gene models from GENCODEv26 (promoters, intronic, exonic, 5’UTR, 3’UTR, intergenic), CpG islands (CGIs) and related features from UCSC (shores, shelves, OpenSea), putative enhancer regions (Enhancers and High Confidence Enhancers from PsychENCODE [18] and H3K27ac [19]). 5- group = CG-DMRs or blocks between all 5 tissue groups. D) As in C, showing enrichment in regions of open chromatin in NeuN- and NeuN+ nuclei and NeuN+ nuclei isolated from the indicated brain regions (PV Cortex = Primary Visual Cortex; Med. Thalamus = Mediodorsal Thalamus) from [10]. E) Example CG-DMRs covering the *NPTXR* gene showing average methylation values for NeuN+ nuclei from each tissue group color coded as in B. Regions of differential methylation are shaded in pink. F) Expression of *NPTXR* from sample matched bulk brain tissues from GTEx v8 data release.

To facilitate interpretation of our data, we conducted a simpler analysis. Specifically, we collapsed frontal cortex and anterior cingulate cortex samples into a “cortical” group and caudate, putamen, and nucleus accumbens into a “basal ganglia”. The resulting 5 tissue groups are consistent with the developmental origins of the brain regions; the telencephalon gives rise to the cerebral cortex (which branches into frontal cortex, anterior cingulate cortex, and the hippocampal formation) and cerebral nuclei (which branches into the amygdala and basal ganglia) while the diencephalon produces the hypothalamus [23]. We identified 181,146 autosomal neuronal CG-DMRs (196 Mb) among these 5 groups covering 11% of all CpGs. Further, the 5-group analysis captured 94% of the CG-DMRs identified in the 8-group analysis (**Supplemental Table S5**). Average DNA methylation levels of the most discriminatory CG-DMRs are, aside from several hippocampus samples, able to segregate samples into their tissue groups (**Fig. 1b**). We also identified 7,671 large regions of differential CpG methylation (which we have previously termed “blocks” of differential CpG methylation; these are identified using a larger bandwidth for smoothing) among the 5 tissue groups (S**upplemental Table S6**). These CG-blocks covered 260 Mb and were on average 33.9 kb in size. CG-DMRs were enriched in enhancer regions identified by PsychENCODE [13], H3K27ac peaks found in the adult brain [24], and in regulatory chromHMM states from 4 brain regions [25] (**Fig. 1c, Fig. S1e**). We also observed enrichment of our CG-DMRs in regions of open chromatin identified in NeuN+ nuclei from 14 brain regions [11] (**Fig. 1d**). Example CG-DMRs within the neuronal pentraxin 1 (NP1) gene (*NPTXR*) are shown (**Fig. 1e**) with hypomethylation in the hippocampal neurons associated with increased expression in bulk hippocampus tissue (**Fig. 1f)**. NP1 is involved in glutamate receptor internalization and has been implicated in Alzheimer’s disease as its upregulation in response to increased amyloid-beta promotes neuronal toxicity [26].

Though we grouped them together in our initial CG-DMR analysis, there is a clear distinction among the basal ganglia tissues. These regions are of particular interest due to their importance in addiction and reward pathways [27], yet no comprehensive analysis of methylation differences in human brain has been performed to date. We performed an additional DMR analysis to assess the methylation differences among neurons from these tissues and identified 16,866 autosomal neuronal basal ganglia CG-DMRs (24 Mb) encompassing 1.7% of all CpGs (**Supplemental Table S7**). Consistent with their regional identity, these basal ganglia CG-DMRs were specifically enriched in open chromatin regions identified in neuronal nuclei from striatal tissues [11] (**Fig. 1d**). Over 13% (2,295/16,866) of these basal ganglia CG-DMRs were not identified in our 5-group CG-DMR analysis. We used the Genomic Regions Enrichment of Annotations Tool (rGREAT v4.0.0) [28] to identify enriched gene ontology terms associated with these unique basal ganglia CG-DMRs. Ten of the top 20 significantly enriched terms were related to ion transport or neuronal signaling (**Supplemental Table S8**). This result suggests that differential methylation near these genes (including Ca^+2^ and K^+^ voltage-gated channel subunit genes) could be involved in fine-tuning their expression in particular neuronal populations within the basal ganglia.

### Differential methylation analysis identifies distinct neuronal subpopulations in the hippocampus

Interestingly, principal component analysis revealed two distinct clusters originating from the hippocampus that were not detected in our previous analysis of hippocampal tissues [12] (**Fig. 1a, Fig. 2a**). The hippocampus is composed of several subregions consisting of four “cornu ammonis” regions and the dentate gyrus. We hypothesized that our samples represented the specific pyramidal and granule neurons within these respective subregions. We tested this hypothesis by identifying autosomal hippocampal CG-DMRs (n = 11,702) between these two clusters (**Fig. 2b, Supplemental Table S9**). GREAT analysis of the top 2,000 hippocampal CG-DMRs showed enrichment in neurogenesis and generation of neurons (**Supplemental Table S8**). As adult neurogenesis occurs in the dentate gyrus, these data suggest that some of these samples originated from that particular subregion. Gene expression data from [29–31], were used to compile a list of 75 genes specifically expressed in dentate gyrus granule neurons (**Supplemental Table S10**) and we intersected hippocampal CG-DMRs with these genes and their promoters (TSS +/-4 kb). We identified 117 hippocampal CG-DMRs overlapping these genes and found that in 12 of the 18 hippocampus samples these marker genes are hypomethylated compared to the other 6 hippocampus samples and the other brain tissues examined (**Fig. 2c**). Specific examination of the *PROX1* gene, a marker of dentate gyrus granule neurons, reveals hypomethylation in the promoter and throughout the gene body in these 12 samples providing strong evidence that these samples were enriched for dentate gyrus neurons (**Fig. 2d**). This group of samples is referred to as dentate gyrus samples throughout the rest of the study bringing the total number of brain regions analyzed to nine.

**Figure 2.**
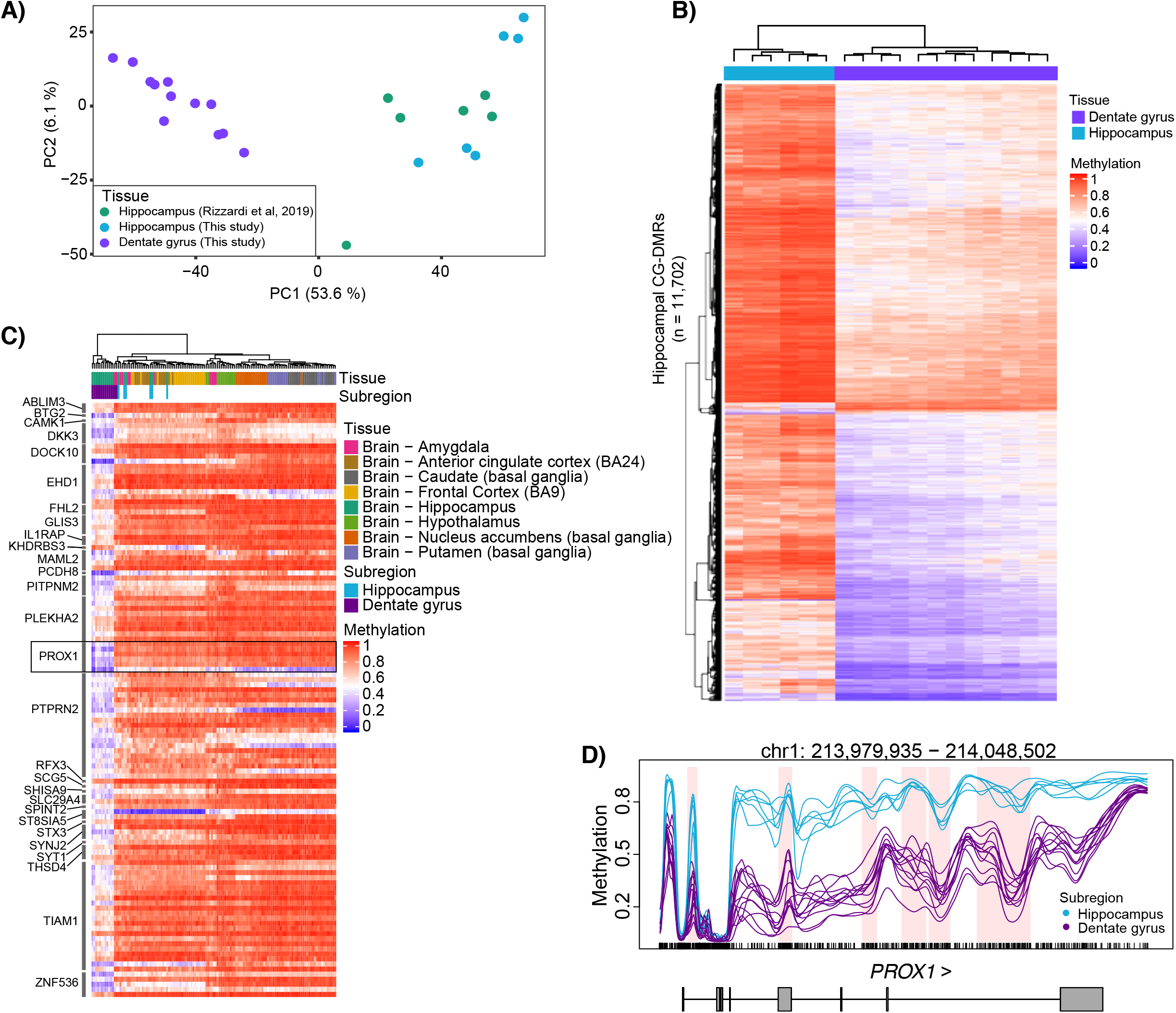
Differential methylation reveals a subset of hippocampus samples originate from the dentate gyrus. A) Principle component analysis of distances derived from average autosomal CpG methylation in 1 kb bins. Data shown are from this study and from [11] as indicated. B) Hierarchical clustering of hippocampus samples based on the average methylation per sample in the CG-DMRs identified between the two hippocampus groups. C) Hierarchical clustering of hippocampus samples based on the average methylation per sample of hippocampal CG-DMRs overlapping dentate gyrus marker genes. The primary marker of dentate gyrus, *PROX1*, is boxed. D) Example hippocampal CG-DMRs in the *PROX1* gene with average methylation values calculated from NeuN+ nuclei isolated from indicated tissue groups and regions of differential methylation shaded pink.

### mCH DMRs

Non-CpG methylation (mCH) is widespread in human neurons and mCH over gene bodies and regulatory elements is generally associated with repression [1, 12, 32]. Interestingly, reduced mCH specifically at neuronal enhancers has recently been associated with Alzheimer’s disease pathology [33]. Paradoxically, mCH has also been associated with lowly transcribed genes involved in neuronal development [34] as well as genes escaping X inactivation [35]. Given the importance of mCH in neuronal development and disease, we performed a differential analysis of mCH across our 5 tissue groups. We identified a total of 264,868 CH-DMRs across all contexts (CA, CT, CC) and strands (+, −) covering a third of the genome (1.0 Gb) (**Supplemental Table S11**). This result represents a >10-fold increase in the number of CH-DMRs identified across the brain compared to our previous work [12]. In that study, we demonstrated high correlations among strands and contexts for mCH, therefore, we use mCA(+) to represent mCH in this study. Global analysis of mCA(+) by principal component analysis revealed segregation of samples based on tissue group though not to the same degree as CpG methylation (**Fig. 3a**). CH-DMRs were 3.5 times broader than CG-DMRs (3,839 vs. 1,086 bp, respectively) and were enriched in CG-DMRs with 67,979 (25%) CH-DMRs overlapping 118,621 CG-DMRs (65%). However, CH-DMRs showed little enrichment for genic or regulatory features and were depleted in CpG islands (**Fig. 3b**). CH-DMRs in the CA(+) context had a median methylation difference of 5.8% with 3,195 having a methylation difference ≥10%. These highly divergent CH-DMRs **(Fig. 3b**; “>10%”) were particularly enriched in genic/intronic and enhancer regions. Results from our CH-DMR analysis among basal ganglia tissues (152,056 CH-DMRs) and between the two hippocampal clusters (100,757 CH-DMRs) were similar to the 5-group CH-DMR analysis (**Supplemental Table S12**). CH-DMRs also showed a slight enrichment in open chromatin across all brain regions analyzed in [11] (**Fig. 3c**). Interestingly, hippocampal and basal ganglia CH-DMRs did not show similar enrichments, but were actually depleted in regions of open chromatin in some tissues. Consistent with CG-DMRs, the highly divergent CH-DMRs were generally hypermethylated in basal ganglia tissues compared to the others (**Fig. 3d)**. CH-DMRs exhibit a high degree of overlap among analyses performed using the 5 tissue groups, basal ganglia samples, and hippocampus samples; this is also true for CG-DMRs (**Fig. 3e, top)**. Additionally, CH-DMRs show substantial overlap with CG-DMRs as we previously reported [12] (**Fig. 3e, bottom**). We can detect many additional CH- and CG-DMRs when looking only among basal ganglia tissues or between hippocampus groups. An example CH-DMR is shown within the gene body of *NRGN*, which encodes the brain-specific protein neurogranin, recently identified as a cerebral spinal fluid biomarker for Alzheimer’s disease [36] (**Fig. 3f**).

**Figure 3.**
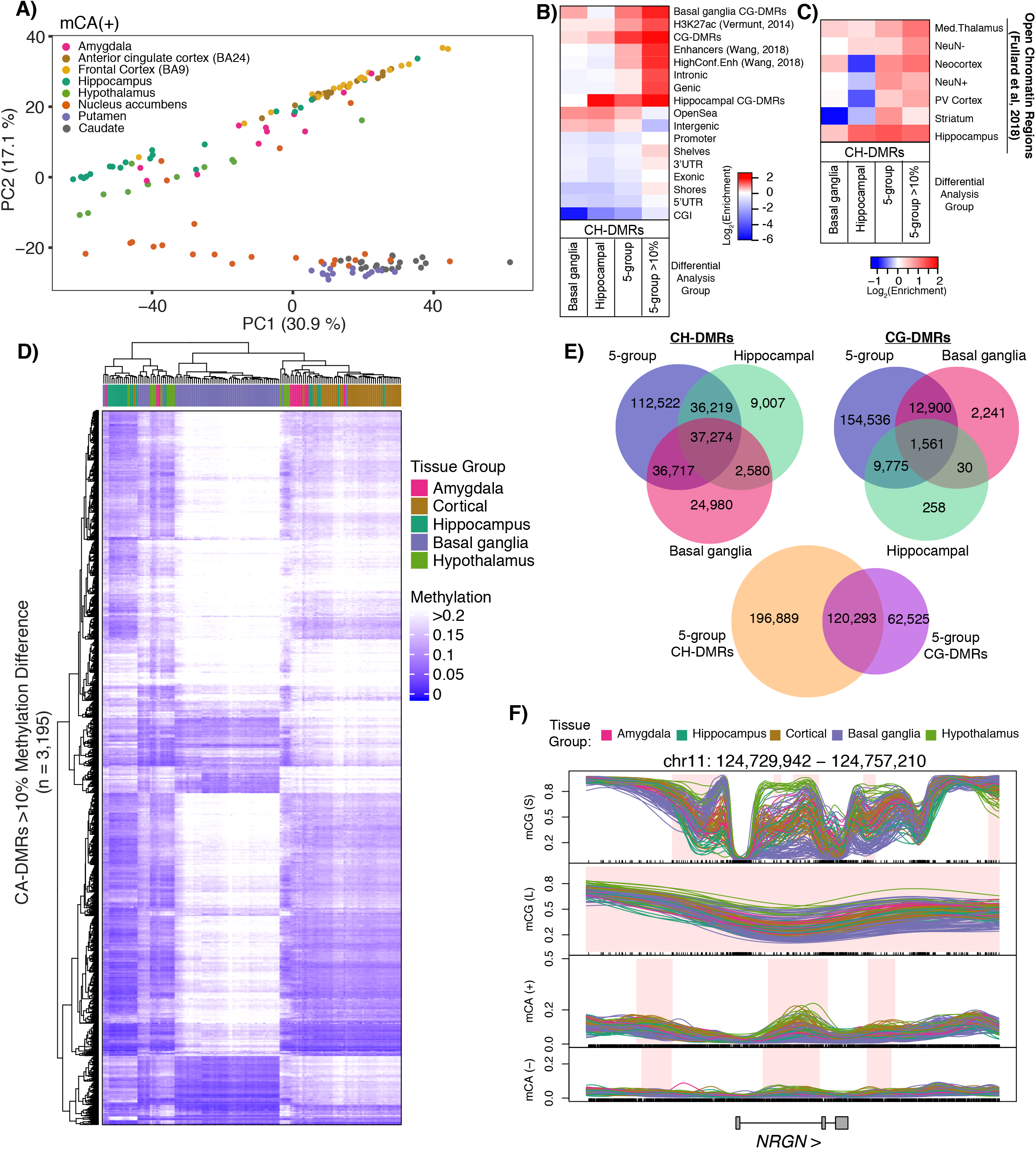
Differential non-CpG methylation across functionally distinct brain regions. **A)** Principle component analysis of distances derived from average autosomal, plus strand CA (mCA+) methylation in 1 kb bins. B) Heatmap representing log2 enrichment of CA, CT, CC within CH-DMRs compared to the rest of the genome for indicated features including CG-DMRs identified in this study. Gene models from GENCODEv26 (promoters, intronic, exonic, 5’UTR, 3’UTR, intergenic), CpG islands (CGIs) and related features from UCSC (shores, shelves, OpenSea), putative enhancer regions (Enhancers and High Confidence Enhancers from PsychENCODE [18] and H3K27ac [19]). 5-group = CH-DMRs between all 5 groups; 5-group >10% = 3,195 CH-DMRs from 5-group comparison with mean CA-DMR methylation difference ≥ 10%. C) As in B, showing enrichment in regions of open chromatin in NeuN- and NeuN+ nuclei and NeuN+ nuclei isolated from the indicated brain regions (PV Cortex = Primary Visual Cortex; Med. Thalamus = Mediodorsal Thalamus) from [10]. D) Hierarchical clustering of samples based on the average CA(+) methylation per sample in the CA-DMRs with >10% methylation difference among the 5 tissue groups. E) Venn diagrams illustrating intersections between CH- and CG-DMRs identified between different analyses. F) Example CA-DMR over *NRGN* with both strands and CG-DMRs (mCG(S); obtained from small smoothing window) and blocks (mCG(L); obtained from large smoothing window). Average methylation values calculated from NeuN+ nuclei isolated from indicated tissue groups. Regions of differential methylation are shaded in pink.

### Identification of VMRs in neurons isolated from human brain tissues

Interindividual variation in DNA methylation has been of interest to many groups and the GTEx sample collection allowed us to explore tissue-specific methylation variability at a genome-wide scale previously not possible. VMRs are loci that are highly variable among individuals within a given tissue type [21, 22]. As a matter of clarification, the word “variability” has been used in other work to refer to changes in DNA methylation between tissues [37], which is not the meaning of VMR used here. Prior studies of methylation variability in brain tissues have been limited to targeted genomic regions (Illumina 450k array) [38] or few individuals [39] using a single brain region. Systemic methylation variability can be driven by genetic effects (methylation QTLs in *cis* and/or *trans*), occur independently as metastable epialleles [40], be caused by environmental exposures, or be confounded by cell type heterogeneity. As we have previously shown [12], much of the variability due to cell type heterogeneity is removed upon isolating NeuN+ nuclei from brain tissues. However, proportions of neuronal subpopulations between brain regions and among individuals could still contribute to variability measurements. Using EpiDISH [41], we estimated the NeuN+ proportions in our samples. For reference data we used sites of differential methylation between NeuN+ and NeuN- nuclei isolated from orbitofrontal cortex identified in Kozlenkov et al [42]. We filtered the 51,412 CpG sites they identified as “neuronal undermethylated” and “glial undermethylated” for |Δβ| > 0.7 resulting in 426 sites. We then eliminated CpGs that overlapped DMRs identified in our 8-group and hippocampal analyses resulting in 201 reference CpGs. This step was critical to eliminate variation due to known region-specific neuronal methylation differences. We found that only four hippocampus samples had any evidence of glial (NeuN-) contamination thus providing independent validation of our sorting efficiency (**Fig. S2a**). Only one of these was less than 97% neuronal with an estimate of 87%.

We identified VMRs by determining the 99^th^ percentile of standard deviation of methylation values in each tissue and applying the lowest standard deviation value (SD = 0.095) as a single cutoff for all tissues (**Fig. S2b**). Using the same SD cutoff allows different tissues to have different numbers of VMRs rather than taking the top most variable regions. This strategy allows for the possibility that some tissues are more variable than others. We identified a total of 81,130 unique VMRs containing > 10 CpGs and covering 159 Mb across all nine brain regions, lung, and thyroid (**Fig. 4a, Fig. S2, Table 1, Supplemental Table S13**). The majority of VMRs are shared among two or more tissues (**Fig. 4b**, “Shared VMR”) with 333 shared among all brain regions. Of those, 202 are “ubiquitous” VMRs, regions of variability shared among all tissues including lung and thyroid (**Fig. 4c**). Remarkably, an average of 24% of the VMRs identified in each tissue are unique to that tissue and we provide examples of tissue-specific VMRs (**Fig.4b)**. To quantify the effect size of the variability, we used the range of the per-sample, across-region average methylation. The median effect size is 35% with almost all VMRs having an effect size greater than 20% and some reaching 50% or higher (**Fig. S2c**).

**Figure 4.**
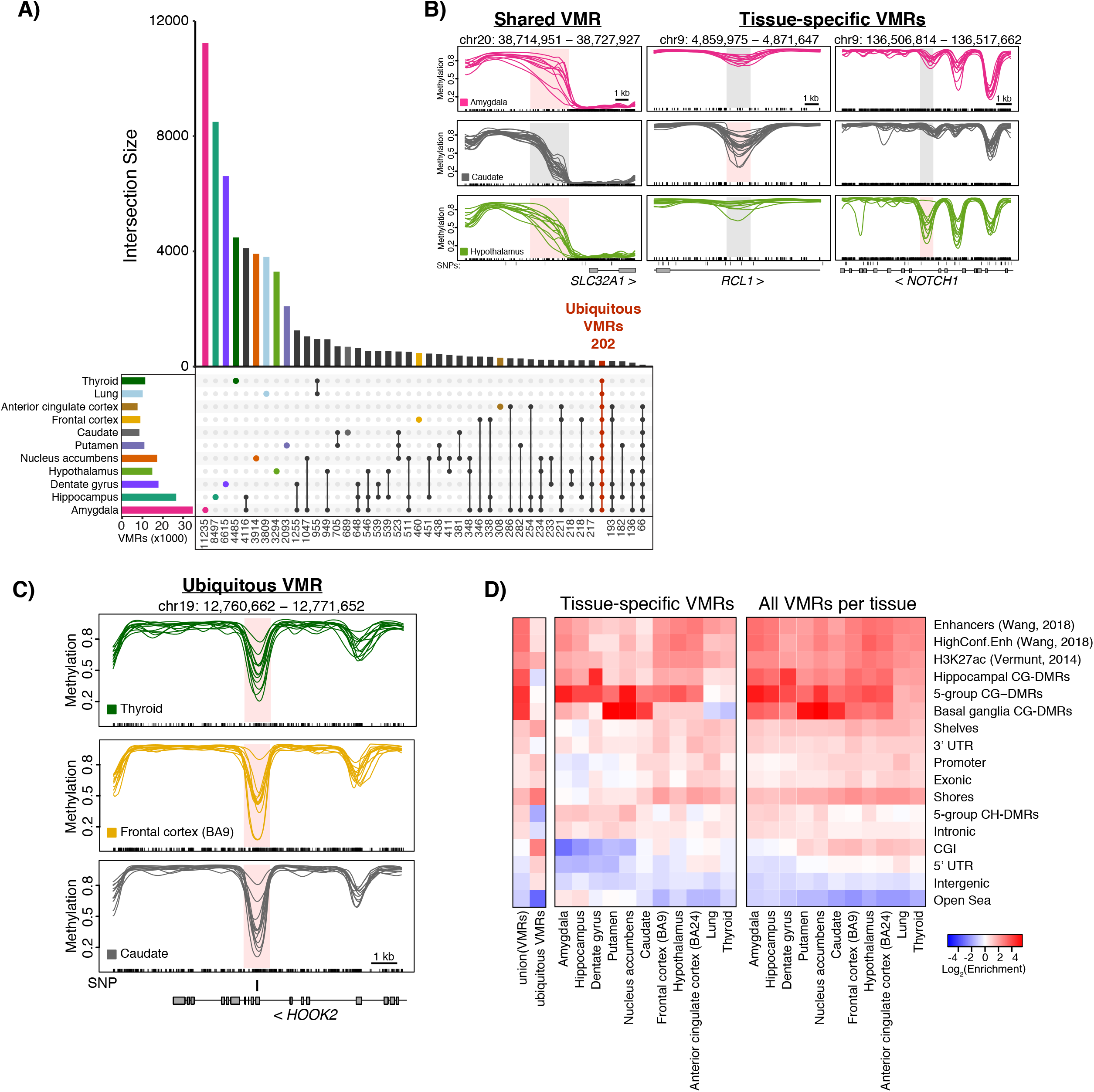
Methylation variability across brain and non-brain tissues. A) VMRs shared among two non-brain tissues and neuronal nuclei from distinct brain regions. Number of VMRs in each intersection is listed at bottom of plot. B) Examples of shared and tissue-specific VMRs in amygdala (pink), caudate (grey), and hypothalamus (green). VMRs are shaded pink in the tissue they were identified in and grey in tissues where they were not considered a VMR; SNPs are indicated when present. C) Example of a ubiquitous VMR shared across brain and non-brain tissues as indicated. D) Heatmap representing log2 enrichment of the union of all VMRs identified, ubiquitous VMRs, tissue-specific VMRs, and all VMRs identified in each tissue as indicated compared to the rest of the genome for genomic features. Gene models from GENCODEv26 (promoters, intronic, exonic, 5’UTR, 3’UTR, intergenic), CpG islands (CGIs) and related features from UCSC (shores, shelves, OpenSea), putative enhancer regions (Enhancers and High Confidence Enhancers from PsychENCODE [18] and H3K27ac [19]). 5-group CG-DMRs = CG-DMRs identified among all 5 tissue groups.

**Table 1.** VMRs

| Tissue | Total |  |  | Tissue-specific |  |  | Overlap<br>CG-DMRs | Within<br>CG-DMRs |
| --- | --- | --- | --- | --- | --- | --- | --- | --- |
|  | VMRs | Mb | CpGs | VMRs | Mb | CpGs |  |  |
| Dentate gyrus | 17,731 | 16.64 | 376,310 | 6,615 | 6.4 | 121,072 | 78.4% | 42.3% |
| Hippocampus | 26,536 | 34.81 | 609,008 | 8,497 | 12.63 | 152,296 | 84.3% | 39.6% |
| Amygdala | 34,876 | 44.98 | 824,870 | 11,235 | 15.77 | 209,042 | 87.9% | 42.9% |
| Putamen | 10,654 | 11.27 | 243,030 | 2,093 | 2.4 | 36,101 | 71.0% | 35.4% |
| Hypothalamus | 14,602 | 14.07 | 342,524 | 3,294 | 3.08 | 59,739 | 76.0% | 42.8% |
| Anterior cingulate cortex | 7,327 | 5.14 | 165,526 | 308 | 0.2 | 4,673 | 76.9% | 50.4% |
| Nucleus accumbens | 17,144 | 18.66 | 392,791 | 3,914 | 4.69 | 69,740 | 84.7% | 46.1% |
| Caudate | 8,137 | 7.19 | 180,073 | 689 | 0.61 | 10,838 | 69.0% | 37.6% |
| Frontal cortex | 8,719 | 6.3 | 205,346 | 460 | 0.25 | 7,804 | 74.5% | 46.6% |
| Lung | 9,770 | 8.59 | 233,724 | 3,809 | 3.29 | 77,481 | 44.1% | 18.6% |
| Thyroid | 11,016 | 10.07 | 258,613 | 4,485 | 3.92 | 86,891 | 44.8% | 19.6% |
| Ubiquitous | 202 | 0.49 | 14,870 |  |  |  | 25.7% | 0.5% |

Almost all (97%) VMRs overlapped a CG-DMR with 35-50% of VMRs fully contained within a CG-DMR (**Table 1**). This percentage drops to 18-20% for VMRs identified in lung and thyroid, which is expected as these two tissues that were not included in CG-DMR analyses. This overlap is reflected in the enrichment of VMRs for CG-DMRs, which is less for lung and thyroid for the reason stated above (**Fig. 4d**). This can be visualized in Fig. 4b (far right panels) where a VMR is present in the hypothalamus (bottom, green) and the mean methylation is significantly different than that in the amygdala (top, pink) or caudate (middle, grey) thus constituting a CG-DMR. These tissue-specific regions of methylation variability were enriched in putative regulatory regions including enhancer- and transcription-associated chromHMM states (**Fig. 4d, Fig. S3a**). VMRs were found across all autosomes (**Fig. S3b**) including the MHC region of chromosome 6 which is known to be highly variable. The MHC region, as well as the pericentromeric region of chromosome 20, harbored more ubiquitous VMRs than any other genomic region similar to previous results [39] **(Fig. S4**). In contrast to most VMRs, ubiquitous VMRs were particularly enriched in CpG islands and shores (**Fig. 4d**).

When considering tissue-specific VMRs, we first focused on the amygdala as 32% of VMRs identified in this tissue were unique to the amygdala. The amygdala displayed the highest interindividual variability among all tissues measured and this was evident both by principal component analysis (**Fig. 1a**) and in the increased number of total VMRs identified (**Table 1**). We hypothesized that, similar to the hippocampus, distinct subregions of the amygdala were isolated from these individuals resulting in increased neuronal heterogeneity. For tissue-specific VMRs, we cannot distinguish true variation from that due to cellular heterogeneity, which led us to investigate how large a contribution heterogeneity makes. Neuroimaging analyses of anatomical and functional connectivity have subdivided the amygdala into as many as 9 distinct subnuclei [43–45]. These subnuclei differ in strength of connectivity to other brain regions including the hypothalamus, hippocampus, and cortical regions. For example, a recent neuroimaging study found that the basolateral nucleus displayed stronger connections to hypothalamus and visual cortex than the centrocortical nucleus which showed stronger connections to the primary motor cortex [46]. As these categorizations are based primarily on neuroimaging data, molecular features of these subregions have yet to be elucidated in the human amygdala. However, single cell RNA-seq of medial amygdala in mouse led to the identification of 16 distinct neuronal subtypes [47]. We examined methylation within 1 kb of the human homologs of the 262 genes used to cluster the neurons and found 649 VMRs, 116 of which were specific to the amygdala. No tissue-specific VMRs from any other brain region were detected near these genes. Hierarchical clustering of amygdala neuronal samples based on these 649 VMRs reveal three groups suggesting that these samples may have originated from distinct subnuclei within the amygdala (**Fig. S5a,b**). VMRs within the *SLC17A7* gene, a marker of glutamatergic neurons, are shown as an example of variable methylation among these three sample groups (**Fig. S5c**). These data strongly suggest that variability among amygdala samples is driven by neuronal subtype differences among the subregions sampled. We were unable to identify VMRs within these three distinct groups as only 3-4 individuals were in each subgroup.

When we consider those VMRs that are not tissue-specific, but are shared among at least one other brain region (as shown in Fig. 4a) it is unlikely those VMRs are due to neuronal heterogeneity, because they are shared between brain regions with distinct neuronal populations. Therefore, the variability of these regions must be shared among different cell types which suggests they have some common biological function. Further analysis of the 949 VMRs shared solely between the amygdala and hypothalamus revealed an enrichment for neurotransmitter transport genes, particularly in the SLC family (**Supplementary Table S14**). There is a VMR ∼3 kb upstream of the TSS of *SLC32A1* (Fig4b, “shared VMR”), which is expressed in GABAergic neurons and mediates uptake of GABA and glycine to synaptic vesicles [48]. These shared VMRs are also found near *SLC6A1* (−15 kb) and *SLC6A11* (+1.4 kb), two other GABA transporters for neurons and glia, respectively, as well as upstream of *SLC6A3* (−52 kb), a dopamine transporter important in pathogenesis of psychiatric disorders [49]. Among the VMRs identified in the other brain regions, 77-95% were shared among two or more. These shared VMRs could be important regions for integrating signaling inputs from neuronal crosstalk.

### DMRs and VMRs are enriched for heritability of brain-linked traits

We and others have shown a strong association between differential epigenetic features and neurological, neuropsychiatric, and behavioral-cognitive phenotypes [4, 11, 12]. Using stratified linkage disequilibrium score regression [50] we asked if the VMRs we identified in each brain region were also associated with brain-linked traits (**Supplemental Table S15)**. First, we replicated our previous findings that regions of differential CpG methylation are enriched for heritability of brain-linked traits including schizophrenia, neuroticism, and depressive symptoms (**Fig. 5a; Fig. S6, Supplemental Table S16).** In addition, CG-DMRs and those identified among basal ganglia regions showed significant enrichment for heritability of attention deficit hyperactivity disorder (ADHD). VMRs identified in hypothalamus were also enriched for heritability of ADHD, while VMRs identified in amygdala, anterior cingulate cortex, and hippocampus were significantly enriched for heritability of schizophrenia (**Fig. 5b)**. Amygdala VMRs showed a greater enrichment than CG-DMRs (6.5 vs 4.6) though they cover ∼75% less of the genome than CG-DMRs (**Supplemental Table S16)**.

**Figure 5.**
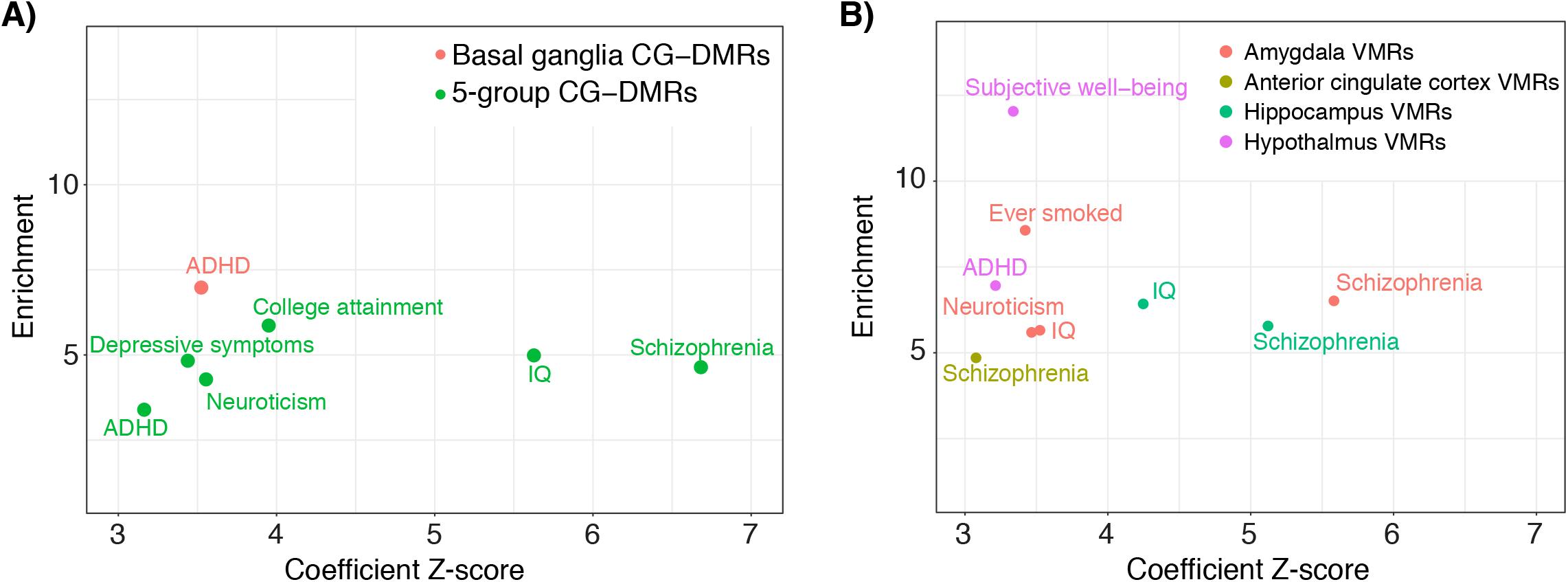
Neuronal CG-DMRs and VMRs are highly enriched for explained heritability of multiple psychiatric, neurological, and behavioral-cognitive traits. Results from running stratified linkage disequilibrium score regression using 30 GWAS traits with 97 baseline features and either DMRs (A) or VMRs (B) identified in this study. Only feature-trait combinations with a coefficient z-score significantly larger than 0 (one-sided z-test with alpha = 0.05, P-values corrected within each trait using Holm’s method) are shown. Enrichment score (y-axis) and coefficient z-score (x-axis) from running this analysis for each of the indicated methylation features combined with the baseline features are plotted.

### Genetic contributions to DNA methylation variability

The most likely explanation for a genetic contribution to DNA methylation variability is a genetic polymorphism inside the VMR region. Roughly 25% of VMRs do not overlap any SNPs with a minor allele frequency (MAF) > 0.1 across our samples (Supplemental Table S17). Genotype data from all but two individuals were available from GTEx v8 [16] leaving 6-20 individuals per tissue, which is too few to conduct a rigorous methylation QTL analysis. However, we did identify several examples of VMRs that overlap one or more SNPs that are associated with altered methylation (**Fig. 6a**,). One example is shown within the *MYO3A* gene which lies upstream of *GAD2*, a primary regulator of GABA synthesis that has been associated with schizophrenia [51], bipolar disorder [52], and major depression [53]. There is at least one *GAD2* enhancer located within the *MYO3A* gene, though it is ∼150 kb away from this VMR [54]. To examine possible genetic contributions to our observed methylation variability, we asked if mQTLs identified from brain tissues were enriched in our VMRs (**Fig. 6b**). The mQTL data we chose were generated using 450k arrays on samples from bulk hippocampus (n = 110) [55], fetal brain (n = 166) [6], and bulk temporal cortex (n = 44) that was also sorted into NeuN+ (n =18) and NeuN- (n = 22) nuclei [56]. We found the greatest enrichments in all datasets within our ubiquitous VMRs consistent with our assertion that methylation variability in this class of VMRs is genetically driven. We also compared our VMRs to two existing datasets [38, 39] that examine genetic and environmental contributions to methylation variation. Garg et al [38] profiled 58 NeuN sorted, frontal cortex samples using the Illumina 450k array and identified 1,136 neuronal VMRs, 996 of which were also detected in our analysis. In addition, they identified 149 VMRs in common among blood, brain, and fibroblast samples and while 142 of these were also identified in our analysis, only 18 were present in our set of ubiquitous VMRs. Gunasekara et al [39] profiled DNA methylation in three tissues from 10 GTEx donors to identify regions where interindividual methylation variation was not tissue-specific. They identified 9,926 correlated regions of systemic interindividual variation (CoRSIVs). We found that our VMRs were enriched for VMRs identified by Garg et al [38] as well as for CoRSIVs [39] (**Fig. 6c**). We detected 16% (1,588/9,926) of CoRSIVs in our analysis. As CoRSIVs are, by definition, consistent across the three tissues sampled (heart, thyroid, and cerebellum), we were unsurprised to find that the enrichment in these regions was lower for tissue-specific VMRs.

**Figure 6.**
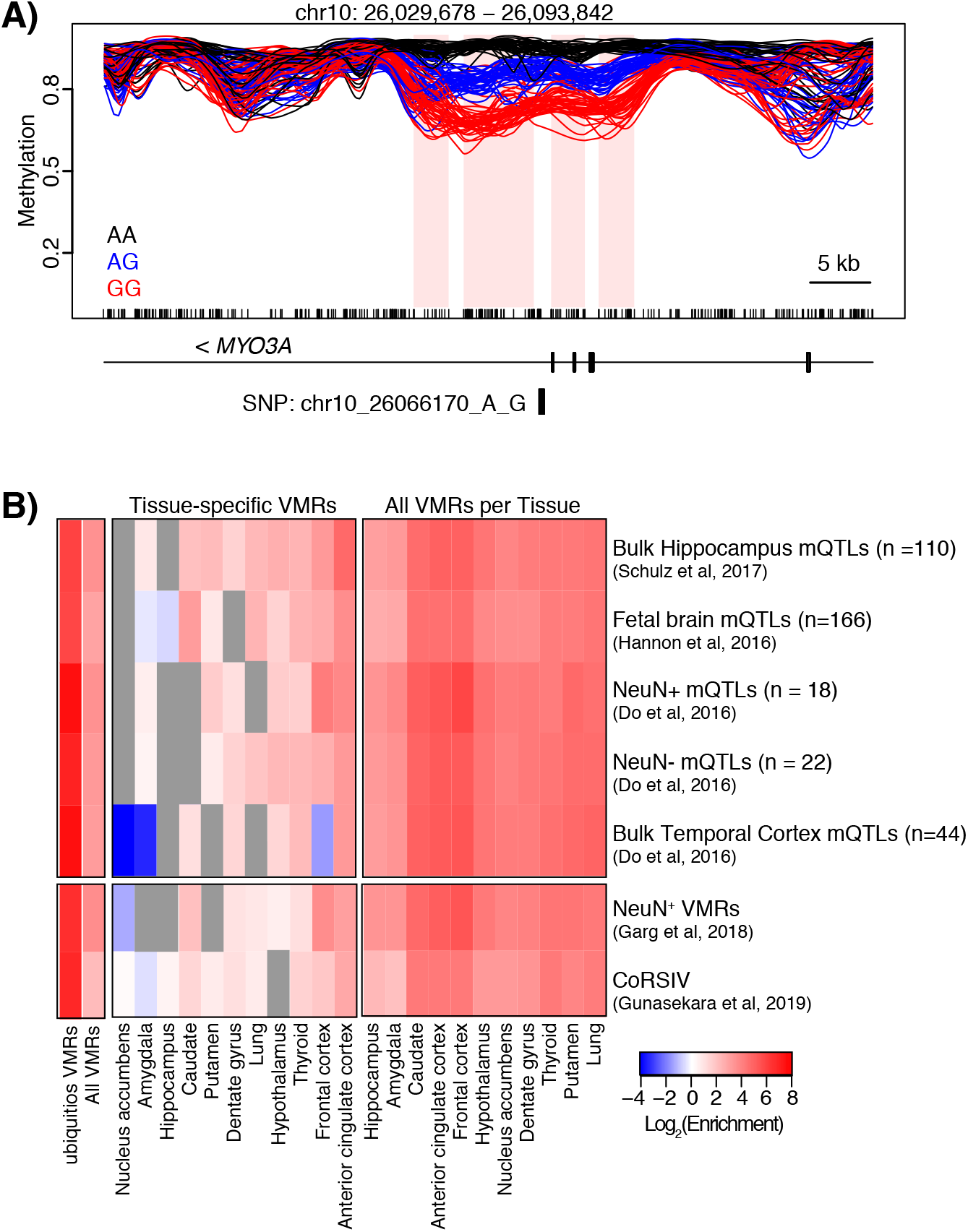
Genetic contributions to methylation variability. A) Example of a SNP associated with altered methylation within *MYO3A* showing average methylation values for each genotype (indicated) for all tissues. The VMRs associated with this SNP are shaded pink and the SNP is indicated. B) Heatmap representing log2 enrichment of CpGs within VMRs and tissue-specific VMRs compared to the rest of the genome for previously published mQTL datasets[6, 55, 56], neuronal VMRs [35] and correlated regions of systemic interindividual variations (CoRSIVs) [36]. Only significant enrichments/depletions are shown; non-significant combinations are shaded grey.

## DISCUSSION

This resource constitutes the largest examination of DNA methylation differences across isolated neurons of the normal adult human brain encompassing 145 samples across 8 brain regions from 12-24 individuals (average of 18). Our focus on neuronal methylation makes this dataset particularly relevant to investigations of neurodegenerative and neuropsychiatric diseases that preferentially impact neuronal function. We have identified 181,146 CG-DMRs and 264,868 CH-DMRs among neurons isolated from amygdala, hypothalamus, hippocampus, cortical and basal ganglia brain regions. This represents a >10-fold increase in both CG- and CH-DMRs compared to our previous study of 4 brain regions in 6 individuals [12]. CG-DMRs were enriched for heritability of brain-related traits including schizophrenia, neuroticism, and depressive symptoms. Importantly, we performed the first analysis of differential methylation among basal ganglia regions (nucleus accumbens, caudate, and putamen) identifying 16,866 neuronal CG-DMRs that were enriched for heritability of ADHD. Interestingly, 2,295 of these CG-DMRs were not identified in the full 5-group analysis and were enriched for genes encoding voltage-gated ion (Ca^2+^, K^+^) channel subunits responsible for maintaining neuronal activity and homeostasis.

Bisulfite sequencing does not distinguish between cytosine methylation and hydroxymethylation. The human brain has elevated levels of hydroxymethylation (∼10-15%) compared to other tissues [57, 58]. In neuronal subpopulations isolated from adult prefrontal cortex, 5hmC levels can be as high as 40% [15], and it will be important in future studies to accurately profile this mark across different brain regions.

Large methylation differences among hippocampus and amygdala neuronal samples enabled the identification of distinct anatomical substructures, particularly the dentate gyrus. We further investigated the differences between the hippocampus and dentate gyrus samples and identified CG-DMRs that were enriched for genes involved in neurogenesis. Substructures within the amygdala are somewhat less well-defined, but we detected three distinct groups based on methylation data that correspond to known features of amygdala neuronal subpopulations identified in mouse [47]. Future studies are needed to fully characterize the substantial neuronal heterogeneity present in the human amygdala.

The large scale of this study enabled us to assess VMRs, regions of inter-individual methylation variability within each brain region genome-wide, including identification of concurrent differences across brain regions in a given individual. Methylation variability can be driven by multiple biological and technical factors including genetic variants, environmental exposures, and cellular heterogeneity. We identified a total of 81,130 VMRs among 9 brain regions, lung, and thyroid. We expected genotype differences to drive variability at shared VMRs and we found that of the 202 ubiquitous VMRs 96% overlapped a SNP (MAF > 0.019). We also identified a substantial number of VMRs that were tissue-specific. The amygdala had the highest number of tissue-specific VMRs and was the most heterogeneous tissue we analyzed suggesting many of these VMRs are driven by cell type differences. Across all tissues, the majority of VMRs (∼75% on average) were shared among two or more brain regions with more overlap among functionally similar tissues (e.g. frontal cortex and anterior cingulate cortex). Methylation variability in this class of VMRs is unlikely to be due to cell type differences within a given tissue as they are shared across multiple distinct neuronal subtypes present in different regions. VMRs shared between the amygdala and hypothalamus were enriched near genes involved in neurotransmitter transport, particularly members of the SLC family of solute carrier transporters. These shared VMRs could represent loci whose methylation is coordinately regulated between interconnected neurons and thus are candidates for further investigation. Importantly, VMRs identified in amygdala, hippocampus, and anterior cingulate cortex were enriched for heritability of schizophrenia suggesting their importance in neuropsychiatric disease. This vast resource of differential methylation and variability will be invaluable to future studies of diverse neuronal functions across the human brain.

## METHODS

### Sample procurement

All tissue specimens and DNA samples were obtained from the GTEx Laboratory Data Analysis and Coordination Center at the Broad Institute. Complete descriptions of the donor enrollment and consent process, and the biospecimen procurement methods, sample fixation, and histopathological review procedures were previously described [19, 59]. Flash-frozen tissues were obtained from the following brain regions: amygdala (n = 12), anterior cingulate cortex (BA24, n = 15), caudate (basal ganglia, n = 22), frontal cortex (BA9, n = 24), hippocampus (n = 20), hypothalamus (n = 13), nucleus accumbens (basal ganglia, n = 23), and putamen (n = 16). Genomic DNA was obtained from lung (n = 18) and thyroid (n = 19). Demographic information for each donor is presented in **Supplementary Table S1**. Summary data and details on data production and processing are also available from the GTEx Portal (http://gtexportal.org).

### Nuclei Extraction, Fluorescence Activated Nuclei Sorting, and DNA Isolation

Total nuclei were isolated as previously described[12] with the following changes. Approximately 100-200 mg of frozen tissue per sample was homogenized in 5 mL of lysis buffer (0.32 M sucrose, 10 mM Tris pH 8.0, 5 mM CaCl_2_, 3 mM Mg acetate, 1 mM DTT, 0.1 mM EDTA, 0.1% Triton X-100) by douncing 50 times in a 40 mL dounce homogenizer. Lysates were transferred to a 18 mL ultracentrifugation tube and 9 mL of sucrose solution (1.8 M sucrose, 10 mM Tris pH 8.0, 3 mM Mg acetate, 1 mM DTT) was dispensed to the bottom of the tube. The samples were then centrifuged at 28,600 rpm for 2 h at 4°C (Beckman Optima XE-90; SW32 Ti rotor). After centrifugation, the supernatant was removed by aspiration and the nuclear pellet was resuspended in 200 uL staining mix (2% normal goat serum, 0.1% BSA, 1:500 anti-NeuN conjugated to AlexaFluor488 (Millipore, cat#: MAB377X) in PBS) and incubated on ice. Unstained nuclei served as the negative control. The fluorescent nuclei were run through a Beckman Coulter MoFlo Cell Sorter with proper gate settings (**Fig. S1**). A small portion of the NeuN^+^ and NeuN^-^ nuclei were re-run on the sorter to validate the purity which was greater than 95%. Only immuno-positive (NeuN^+^) nuclei were collected. DNA was extracted directly from the sorted nuclei without centrifugation using the MasterPure DNA Extraction kit (Epicentre, Madison, Wisconsin, USA) following the manufacturer’s instructions.

### Whole genome bisulfite sequencing (WGBS)

WGBS single indexed libraries were generated using the NEBNext Ultra DNA library Prep kit for Illumina (New England BioLabs, Ipswich, MA, USA) on the Agilent Bravo automated liquid handling platform with a custom high-throughput protocol. 40–300 ng gDNA was quantified by Qubit dsDNA BR assay (Invitrogen, Carlsbad, CA, USA) and 1% unmethylated lambda DNA (cat#: D1521, Promega, Madison, WI, USA) was spiked in to measure bisulfite conversion efficiency. DNA was fragmented to an average insert size of 400-500 bp using the Covaris LE220 Focused-ultrasonicator in a 55 μl volume. The fragmented gDNA was converted to end-repaired, adenylated DNA using the NEBNext Ultra End Repair/dA-Tailing Module (cat#: 7442L, New England BioLabs, Ipswich, MA, USA). Methylated adaptors (NEBNext Multiplex Oligos for Illumina; cat#: E7535L New England BioLabs, Ipswich, MA, USA) were ligated to the product from the preceding step using the NEBNext Ultra Ligation Module (cat#: 7445L, New England BioLabs, Ipswich, MA, USA). Size-selection was performed using AMPure XP beads and insert sizes of ∼400 bp were isolated (0.37x and 0.20x ratios). Samples were bisulfite converted after size selection using the EZ-96 DNA Methylation-Gold Kit (cat#: D5008, Zymo, Irvine, CA, USA) following the manufacturer’s instructions. Amplification was performed following bisulfite conversion using primers from the NEBNext Multiplex Oligos for Illumina module (cat#: E7535L, New England BioLabs, Ipswich, MA, USA) and the Kapa HiFi Uracil+ PCR system (cat#: KK2801, Kapa Biosystems, Boston, MA, USA) with the following cycling parameters: 98°C 45 sec / 8 cycles: 98°C 15 sec, 65°C 30 sec, 72°C 30 sec / 72°C 1 min. The PCR enriched product was cleaned up using 0.9x AMPure XP beads (cat#: A63881, Beckman Coulter, Brea, CA, USA). Final libraries were run on 2100 Bioanalyzer (Agilent, Santa Clara, CA, USA) using the High-Sensitivity DNA assay; samples were also run on Bioanalyzer after shearing and size selection for quality control purposes. Libraries were quantified by qPCR using the Library Quantification Kit for Illumina sequencing platforms (cat#: KK4824, KAPA Biosystems, Boston, MA, USA), using 7900HT Real Time PCR System (Applied Biosystems). Libraries were sequenced with the Illumina HiSeq4000 using 151 bp paired-end run with a 27% PhiX spike-in.

### Mapping and quality control of WGBS reads

We trimmed reads of their adapter sequences using Trim Galore! (v0.4.0) (http://www.bioinformatics.babraham.ac.uk/projects/trim_galore/) and quality-trimmed using a cutoff of 30. We then aligned these trimmed reads to the hg38 build of the human genome [including autosomes, sex chromosomes, mitochondrial sequence (available from https://software.broadinstitute.org/gatk/download/bundle) plus lambda phage (accession NC_001416.1) but excluding non-chromosomal sequences] using Bismark[60] (v0.19.0) with the following alignment parameters: bismark --bowtie2 −1 ${READ1} −2 ${READ2}. **Supplementary Table S2** summarizes the alignment results. Using the reads aligned to the lambda phage genome, we estimated that all libraries had a bisulfite conversion rate > 99%.

We then used bismark_methylation_extractor to summarize the number of reads supporting a methylated cytosine and the number of reads supported a unmethylated cytosine for every cytosine in the reference genome. Specifically, we first computed and visually inspected the M-bias[61] of our libraries. Based on these results, we decided to ignore the first 2 bp and last 1 bp of read1 and the first and last 3 bp of read2 in the subsequent call to bismark_methylation_extractor with parameters: --ignore 2 --ignore_r2 3 --ignore_3prime 1 -- ignore_3prime_r2 3. The final cytosine report file summarizes the methylation evidence at each cytosine in the reference genome.

To confirm the genotypes of our samples matched the correct individual GTEx donors, we first downloaded the GTEx v8 genotype data from dbGaP. We validated our genotypes with 58 SNPs used for quality control with the MethylationEPIC BeadChip microarray (Illumina) [62]. For each aligned bam file we created a vcf of these SNPs using: samtools mpileup -v -t DP,AD -f hg38.fasta –-positions 58_SNPs.bed -R -b list_of_bamfiles.txt | bcftools call -m > sample.vcf. The resulting vcf files were indexed and used as input to the bcftools gtcheck command. If the participant ID with the lowest discordance value did not match the sample participant ID then the sample was removed from the analysis as failing genotype QC. Thirty samples failed this quality control measure.

### EpiDISH Analysis

To independently validate our sorting efficiency, we used the robust partial correlations (RPC) method from the EpiDISH R pacakge v2.4.0 [41] to estimate the proportion of non-neuronal nuclei in our NeuN+ samples. The reference sites used were the 23,670 “neuronal undermethylated” and 27,742 “glial undermethylated” CpGs identified in [42]. We filtered for |Δβ| > 0.7 resulting in 426 sites. To remove any known sites of region-specific methylation variation, we removed an addition 209 CpGs that overlapped our 8-group and hippocampal DMRs for a final reference set of 201 CpGs. Neuronal proportions were estimated from smoothed BS β-values from all our NeuN+ samples.

### Annotation and External Data

The hg38 build of the human reference genome was used for all analyses. Genes, exons, introns, and UTRs were taken from GENCODE v26 (https://www.gencodegenes.org/human/release_26.html)[63]. Gene bodies were defined by taking the union over all transcripts (transcription start site to transcription end site) for each gene. Promoters were defined as 4 kb centered on the transcription start site. CpG islands were downloaded from UCSC (http://genome.ucsc.edu/) [64, 65]. CpG shores are defined as 2 kb flanking CpG islands and CpG shelves are defined as 2 kb flanking CpG islands. The 15-state chromHMM model for 7 adult brain tissues from the Roadmap Epigenomics Project[25] was downloaded using the R/Bioconductor AnnotationHub package (v2.6.4). The selected brain regions and their Roadmap Epigenomics codes were: Brain Angular Gyrus (E067); Brain Anterior Caudate (E068); Brain Cingulate Gyrus (E069); Brain Germinal Matrix (E070); Brain Hippocampus Middle (E071); Brain Inferior Temporal Lobe (E072); Brain Dorsolateral Prefrontal Cortex (E073); Brain Substantia Nigra (E074); Fetal Brain Male (E081); Fetal Brain Female (E082). Separate enrichments were calculated for each chromHMM annotation then averaged together for graphical representation. The PsychENCODE enhancer set for the prefrontal cortex was taken from [13] (http://resource.psychencode.org/). Brain enhancers denoted by H3K27ac were obtained from [24]. Regions of open chromatin in neuronal and non-neuronal nuclei from 14 brain regions were obtained from Table S4 from [11].

For the analysis of hippocampal subgroups, we used gene expression data from [29–31] to compile a list of 75 genes specifically expressed in dentate gyrus granule neurons (**Supplementary Table S10**). We examined DNA methylation of the hippocampal CG-DMRs that overlapped these genes or their promoters (TSS +/- 4 kb). For the analysis of amygdala subregions, we profiled DNA methylation within 1 kb of the TSS of 216 genes used to cluster single cell expression data from mouse medial amygdala neurons (Supplementary Table S2 in [47]). We first converted the mouse genes to their human homologs and position in hg38 using the biomaRt v2.38.0 [66, 67] R package. We examined DNA methylation at VMRs identified in the amygdala that were within 1 kb of these genes. Any data originally mapped to hg19 were lifted over to hg38 using liftOver from the rtracklayer R package [68].

Gene expression data was downloaded from GTEx v8 (dbGaP Accession: phs000424.v8.p2). While these expression data were generated from the same donors and tissues used in our study, gene expression was measured in bulk tissues rather than in sorted neuronal nuclei. Therefore, no large-scale integrative analyses were performed with these data.

### DMR analyses

Differentially methylated regions and blocks of differential methylation were identified and annotated as previously described [12] except a minimum of 70 CpGs were required in a smoothing window. Large blocks of differential methylation were identified after smoothing over windows of at least 20 kb containing at least 500 CpGs. Following smoothing, we analyzed all CpGs that had a sequencing coverage of at least 1 in all samples for a total of 27,660,298 CpGs. Unlike CpGs, CpAs and CpTs are not palindromic, so were analyzed separately for each strand, for a total of 4 strand/dinucleotide combinations:

- mCA (forward strand)
- mCA (reverse strand)
- mCT (forward strand)
- mCT (reverse strand)

For each dinucleotide/strand combination we ran a single ‘small-ish’ smooth to identify DMRs (smoothing over windows of at least 3 kb containing at least 200 CpAs or CpTs). Following smoothing, we analyzed all CpAs and CpTs regardless of sequencing coverage. We performed five distinct CpG and non-CpG DMR analyses. The first analysis identified both CpG and non-CpG (CH) DMRs among neuronal nuclei isolated from 5 tissue groups: cortical (BA9, BA24), basal ganglia (putamen, nucleus accumbens, caudate), hippocampus, hypothalamus, and amygdala brain regions. To identify the most discriminatory CG-DMRs (in **Fig. 1b**), we used the annotations to identify those DMRs unique to a specific group. For example, DMRs identified only when amygdala was compared to the other four groups, and not when those four groups were compared to each other. This resulted in a total of 27,782 discriminatory CG-DMRs:

1. Basal ganglia CG-DMRs = 10,000 (sorted by maxStat and took the top 10,000)
2. Cortical CG-DMRs = 1,604
3. Amygdala CG-DMRs = 14
4. Hippocampus CG-DMRs = 14,158
5. Hypothalamus CG-DMRs = 2,006

We also performed the same analyses across all 8 tissues individually. We next looked for DMRs among neuronal nuclei within the basal ganglia tissues. Finally, we identified DMRs among the two groups of hippocampus samples to determine why they were segregating into two distinct clusters identified by principal component analysis. We ordered these hippocampal CG-DMRs by absolute value of areaStat and used the top 2,000 DMRs for analysis of gene ontology using rGREAT v4.0.0 [28].

### VMR analyses

For VMR analysis, we considered 24,630,044 CpGs that had sequencing coverage of ≥ 5 reads in ≥ 100 samples. To identify variably methylated regions (VMRs), genome-wide standard deviations of DNA methylation were determined and an standard deviation of 0.095 was chosen as our cutoff for all tissues ensuring that at least 99% of the methylation distribution is within +/- 2 standard deviations (equivalent to 38% methylation difference). VMRs were then filtered to include those with >10 CpGs and with a Cook’s distance > 0.7 to remove VMRs driven by sample outliers.

### Enrichment of DMRs, blocks, and VMRs in genomic features

We formed a 2×2 contingency table of (n_11_, n_12_, n_21_, n_22_); specific values of (n_11_, n_12_, n_21_, n_22_) are described below. The enrichment log odds ratio was estimated by log_2_(OR) = log_2_(n_11_) + log_2_(n_22_) - log_2_(n_12_) - log_2_(n_21_), its standard error was estimated by se(log2(OR)) = sqrt(1 / n_11_ + 1 / n_12_ + 1 / n_21_ + 1 / n_22_), and an approximate 95% confidence interval formed by [log_2_(OR) – 2 × se(log2(OR)), log_2_(OR) + 2 × se(log2(OR))]. As the odds ratio is equivalent to enrichment, for clarity figures show log_2_(enrichment) rather than log_2_(OR). We also report the P-value obtained from performing Fisher’s exact test for testing the null of independence of rows and columns in the 2×2 table (i.e. the null of no enrichment or depletion) using the fisher.test() function from the ‘stats’ package in R [69].

We computed the enrichment of CpXs (CpGs, CpAs, or CpTs, as appropriate) within DMRs, blocks, or VMRs inside each genomic feature (e.g., exons, enhancers, etc.). Specifically, for each genomic feature, we constructed the 2×2 table (n_11_, n_12_, n_21_, n_22_), where:

- n_11_ = Number of CpXs in DMRs/blocks/VMRs that were inside the feature
- n_12_ = Number of CpXs in DMRs/blocks/VMRs that were outside the feature
- n_21_ = Number of CpXs not in DMRs/blocks/VMRs that were inside the feature
- n_22_ = Number of CpXs not in DMRs/blocks/VMRs that were outside the feature

The total number of CpXs, n = n_11_ + n_12_ + n_21_ + n_22_, was the number of autosomal CpXs in the reference genome. We counted CpXs rather than the number of DMRs or bases because this accounts for the non-uniform distribution of CpXs along the genome and avoids double-counting DMRs that are both inside and outside the feature.

### Genotype Analysis

We utilized genotypes from GTEx Analysis Release v8 (dbGaP Accession: phs000424.v8.p2) to identify SNPs associated with methylation values at VMRs. We first removed SNPs in linkage disequilibrium, removed missing genotypes, and kept only autosomal SNPs using Plink v2.0 (www.cog-genomics.org/plink/2.0/) [70] with the following parameters: `- -geno 0.1 --indep-pairwise 100’kb’ 1 0.2 --autosome`. We then used Plink to filter the dataset to include only those 26 individuals for which we generated DNA methylation data and retained only SNPs with MAF ≥ 0.1 resulting in 149,185 autosomal SNPs. A SNP matrix was created from the resulting vcf using the readVCF and genotypeToSnpMatrix functions in the VariantAnnotation (v1.28.13) [71] R package.

### Stratified linkage disequilibrium score regression

We used stratified linkage disequilibrium score regression (SLDSR), implemented in the LDSC [72] software, to evaluate the enrichment of common genetic variants from genome-wide association study (GWAS) signals to partition trait heritability within functional categories represented by our DMRs and VMRs. SLDSR estimates the proportion of genome-wide single nucleotide polymorphism (SNP)-based heritability that can be attributed to SNPs within a given genomic feature by a regression model that combines GWAS summary statistics with estimates of linkage disequilibrium from an ancestry-matched reference panel. Links to GWAS summary statistics are available in **Supplementary Table S15**. The GRCh38 “baseline-LD model v2.2” data files were downloaded from https://data.broadinstitute.org/alkesgroup/LDSCORE/ following instructions at https://github.com/bulik/ldsc/wiki.

We ran LDSC (v1.0.0; https://github.com/bulik/ldsc) to estimate the proportion of genome-wide SNP-based heritability of 30 traits (**Supplementary Table S15**) across the 97 ‘baseline’ genomic features and our neuronal (NeuN+) DMRs and VMRs:

1. 5-group CG-DMRs: CG-DMRs among 5 brain tissue groups (196 Mb) (**Supplementary Table S5**)
2. 5-group CH-DMRs: Union of CA-, CC-, and CT-DMRs (both strands) among 5 brain tissue groups (1,010 Mb) (**Supplementary Table S11**)
3. Basal ganglia CG-DMRs: CG-DMRs among 3 basal ganglia tissues (24 Mb) (**Supplementary Table S7**)
4. Basal ganglia CH-DMRs: Union of CA-, CC-, and CT-DMRs (both strands) among 3 basal ganglia tissues (284.4 Mb) (**Supplementary Table S12**)
5. Hippocampal CG-DMRs: CG-DMRs among the two hippocampal groups (24.4 Mb) (**Supplementary Table S9**)
6. Hippocampal CH-DMRs: Union of CA-, CC-, and CT-DMRs (both strands) among the two hippocampal groups (596 Mb) (**Supplementary Table S12**)
7. VMRs identified in each brain tissue (Mb covered listed in Table 1) (**Supplementary Table S13**)

We performed a standard SLDSR analysis, as suggested by the method authors, whereby each of the brain-specific features was added one at a time to a ‘full baseline model’ that included the 97 ‘baseline’ categories that capture a broad set of genomic annotations. We used SLDSR to estimate a ‘coefficient z-score’ and an ‘enrichment score’ for each feature-trait combination. A brief description of their interpretation is given below; we refer the interested reader to the Online Methods of [50] for the complete mathematical derivation. A coefficient z-score statistically larger than zero indicates that adding the feature to the model increased the explained heritability of the trait, beyond the heritability explained by other features in the model. The enrichment score is defined as (proportion of heritability explained by the feature) / (proportion of SNPs in the feature). The enrichment score is unadjusted for the other features in the model, but is more readily interpretable as an effect size. Particularly interesting are those feature-trait combinations with statistically significant z-score coefficients and large enrichment scores. Z-score coefficient p-values within each trait were post-hoc adjusted for multiple testing using Holm’s method [73] (**Supplementary Table S16**).

### Supplementary Software

All statistical analyses were performed using R [69] (v3.5.x) and made use of packages contributed to the Bioconductor project [66, 74]. In addition to those R/Bioconductor packages specifically referenced in the above, we made use of several other packages in preparing results for the manuscript:

- bsseq (v1.14)
- AnnotationHub (v2.6.4)
- biomaRt [66, 67] (v2.30.0)
- GenomicAlignments [75] (v1.10.0)
- GenomicFeatures [75] (v1.26.2)
- GenomicRanges [75] (v1.26.2)
- ggplot2 [76] (v2.2.1)
- Hmisc (v4.0-2)
- Matrix (v1.2-8)
- Rtracklayer [68] (v1.34.1)
- SummarizedExperiment (v1.4.0)
- EnrichedHeatmap (v1.4.0)
- Picard (v2.2.2)
- seqtk (v1.2-r94)
- tximport (v1.2.0)
- rGREAT (v4.0.0)

### Availability of data and materials

All GTEx open-access data are available on the GTEx Portal (https://gtexportal.org/home/datasets). The genotype data used for the analyses described in this manuscript were obtained from dbGaP accession number phs000424.v8.p2 on 12/13/2017. All GTEx protected data are available via application to dbGaP (**accession phs000424**). Access to the raw sequence data is now provided through the AnVIL platform (https://gtexportal.org/home/protectedDataAccess). The code used for all analyses in this manuscript is available at https://github.com/hansenlab/egtex_brain_wgbs. Data are made available as a UCSC track hub that can be accessed by going to https://genome.ucsc.edu/cgi-bin/hgHubConnect#unlistedHubs and loading https://egtex-wgbs.s3.amazonaws.com/hub.txt.

## Acknowledgements

We thank the donors and their families for their generous gifts of organ donation for transplantation, and tissue donations for the GTEx research project.

## Author Contributions

L.F.R, K.D.H, and A.P.F designed the study; L.F.R. and H.Z. performed nuclei sorting with the assistance of K.E.S. and S.D.T.; R.T., A.I., C.M.C., L.F.R. performed DNA extractions; A.I, R.T., C.M.C. performed WGBS library preparation and sequencing; S.R. and L.F.R. established sequence processing pipeline, A.P.F. oversaw the experiments; K.D.H. oversaw the data analysis. L.F.R, P.F.H., and K.D.H performed data analysis; L.F.R, P.F.H., K.D.H, and A.P.F. interpreted the results; L.F.R, K.D.H, and A.P.F wrote manuscript. All authors read and approved the final manuscript.

## Additional Materials

### Supplementary Figures

**Figure S1.**
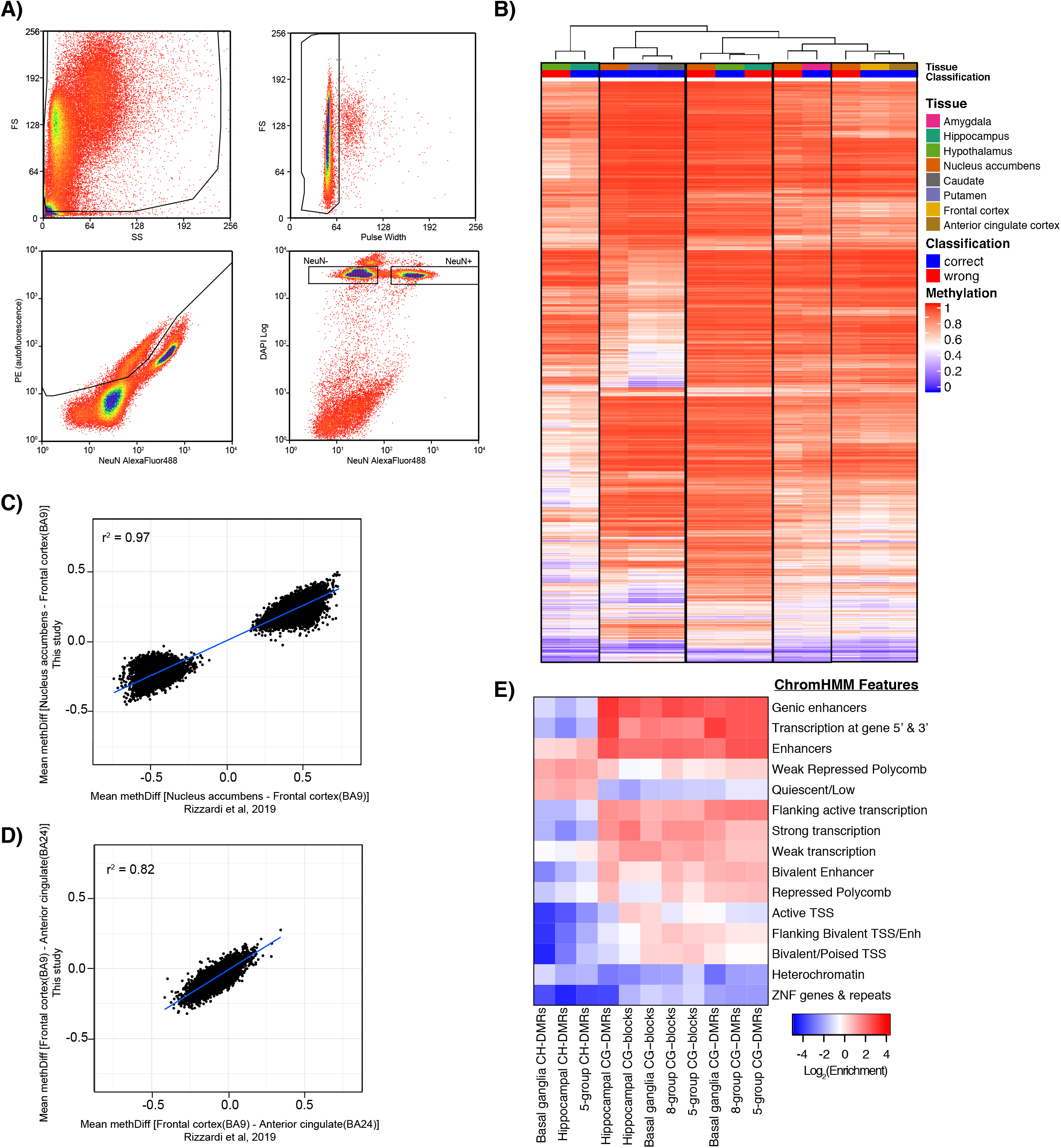
Isolation and characterization of neuronal (NeuN+) nuclei from frozen brain tissue samples. (A) Nuclei in this representative example from caudate tissue were isolated by fluorescence activated nuclei sorting from caudate and debris, doublets, and auto-fluorescent nuclei were gated out as shown. The remaining nuclei were separated based on detection of AlexaFluor 488-conjugated anti-NeuN antibody (MAB377X). (B) Several samples were outliers based on principal component analysis and were eliminated from further analysis. To confirm that they were incorrectly labeled, we averaged the methylation values from all other samples in each tissue group over the CG-DMRs identified among the 5 tissue groups. We then performed hierarchical clustering of the outlier samples (red) along with the merged tissue groups (blue). This analysis revealed the most likely tissue of origin for these outlier samples and confirmed their incorrect labeling. C) Comparison of methylation differences at DMRs identified in [12] between nucleus accumbens and frontal cortex (BA9) NeuN+ samples from [12] and this study. D) Comparison of methylation differences between frontal cortex (BA9) and anterior cingulate cortex (BA24) NeuN+ samples from [12] (x-axis) and this study (y-axis) at all 13,074 DMRs identified in [12]. E) Heatmap representing mean log2 enrichment of all CG-DMRs, CG-blocks, and CH-DMRs identified among the 5 tissue groups as indicated compared to the rest of the genome for chromHMM states from 4 brain regions (15-state model).

**Figure S2.**
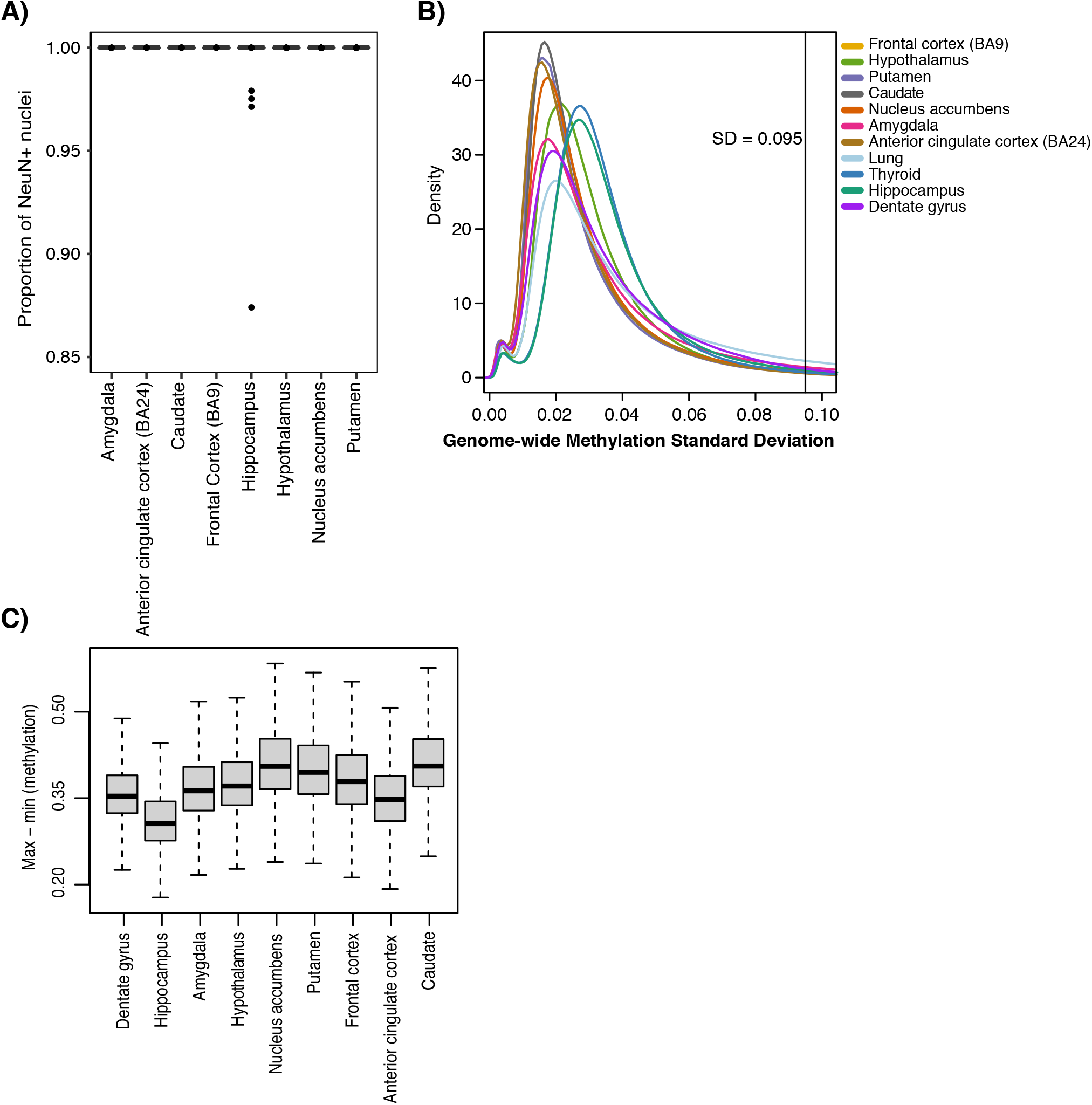
Validation of neuronal proportions and variability measurements. A) Estimates of neuronal proportions using EpiDISH. B) Distribution of standard deviation values within each tissue with the VMR cutoff value indicated. C) The distribution of VMR effect sizes, defined as maximum – minimum of the per-region, sample specific methylation.

**Figure S3.**
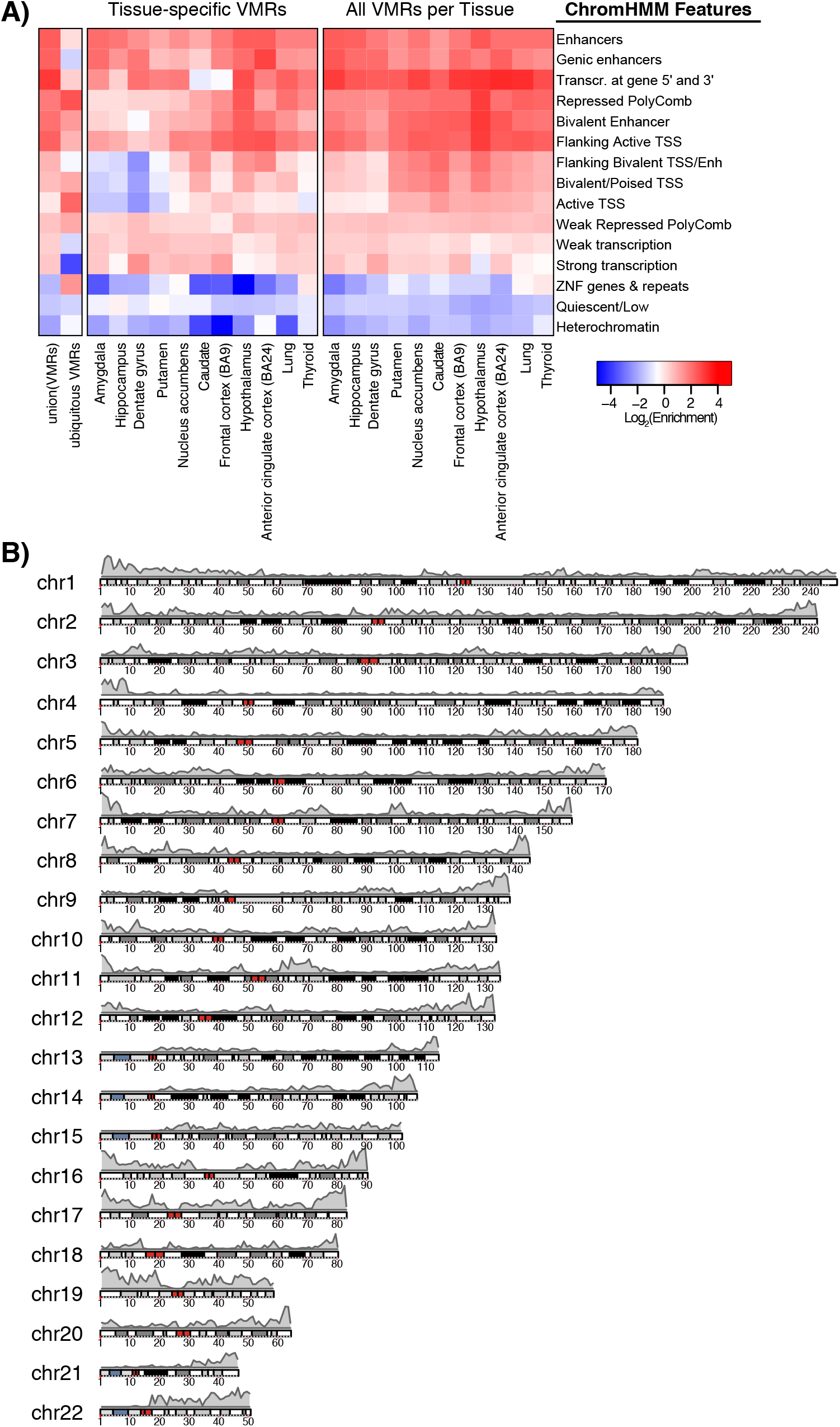
ChromHMM enrichment and distribution of VMRs. A) Heatmap representing log2 enrichment of the union of all VMRs identified. ubiquitous VMRs, tissue-specific VMRs, and all VMRs identified in each tissue as indicated compared to the rest of the genome for chromHMM states from 4 brain regions (15-state model). B) Distribution of all VMRs across the genome.

**Figure S4.**
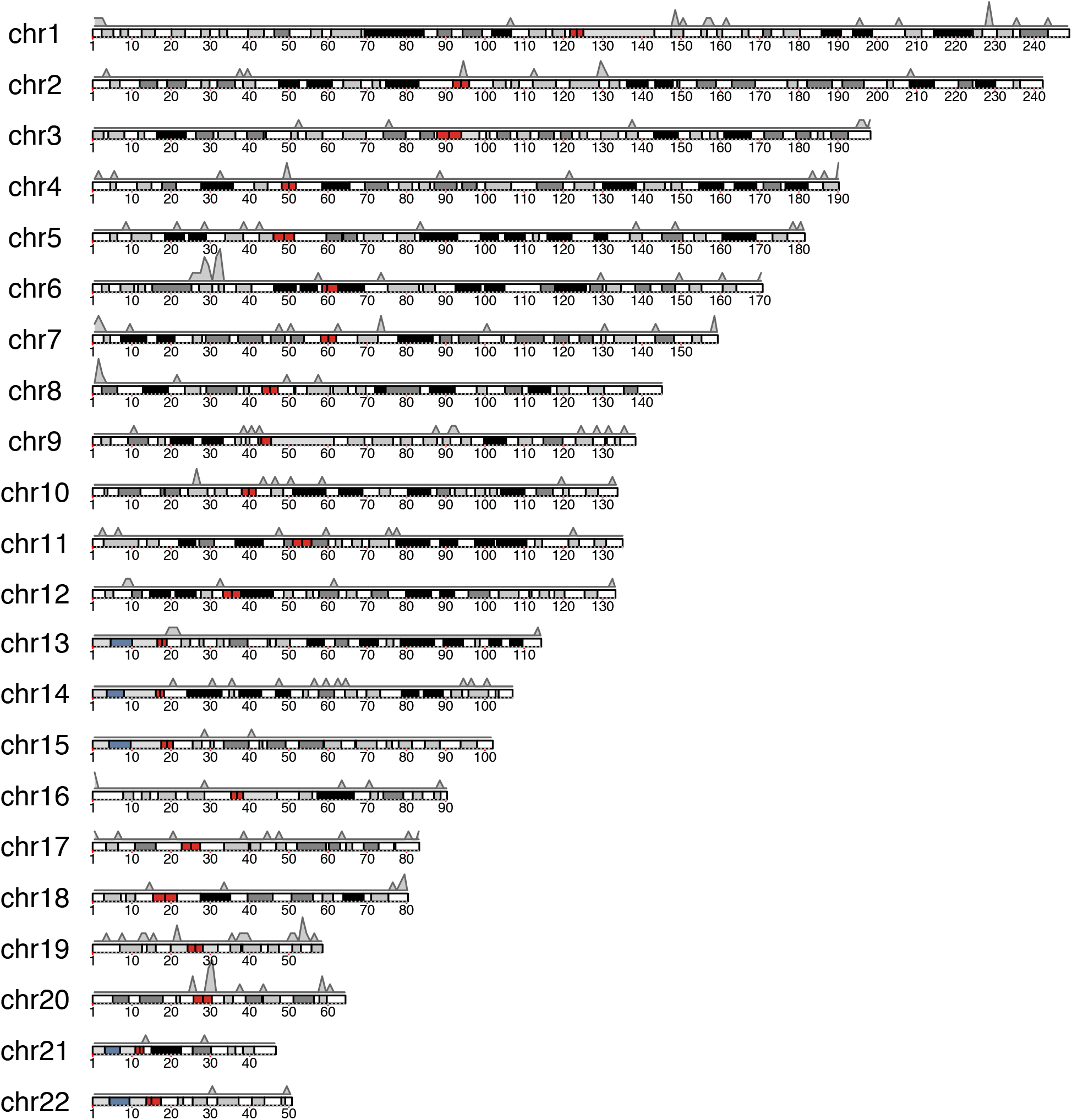
Distribution of ubiquitous VMRs across the genome.

**Figure S5.**
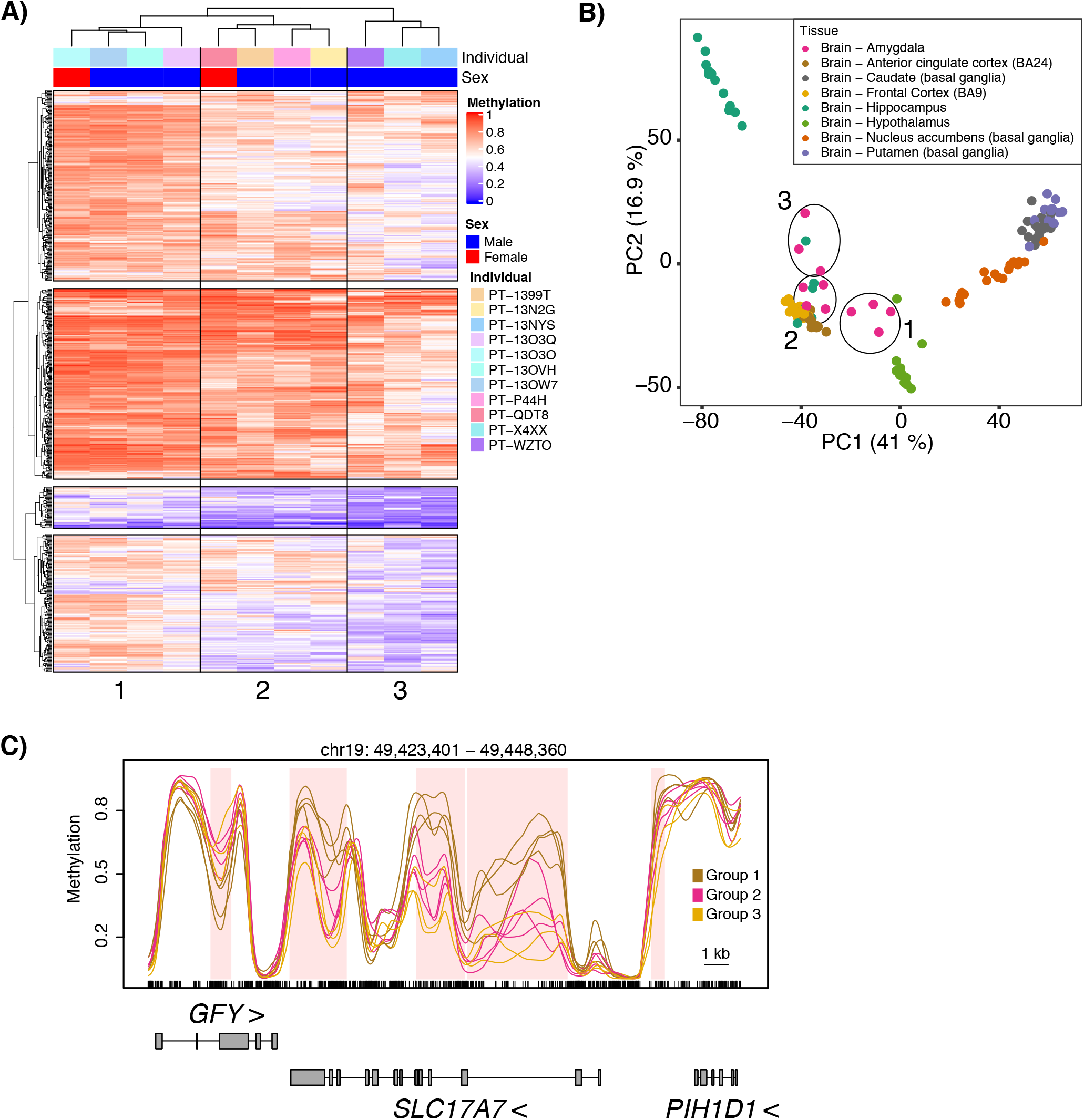
Neuronal heterogeneity among subregions of the amygdala contribute to methylation variability. (A) Hierarchical clustering of amygdala samples based on the average methylation in the 649 VMRs identified in amygdala within 1 kb of the human homologs of genes that identify distinct subregions within the amygdala. (B) Identification of the three amygdala subgroups on the initial principal component analysis from Figure 1A. (C) Example VMRs within the *SLC17A7* gene showing average methylation values for each amygdala subgroup identified in B as indicated. VMRs are shaded pink.

**Figure S6.**
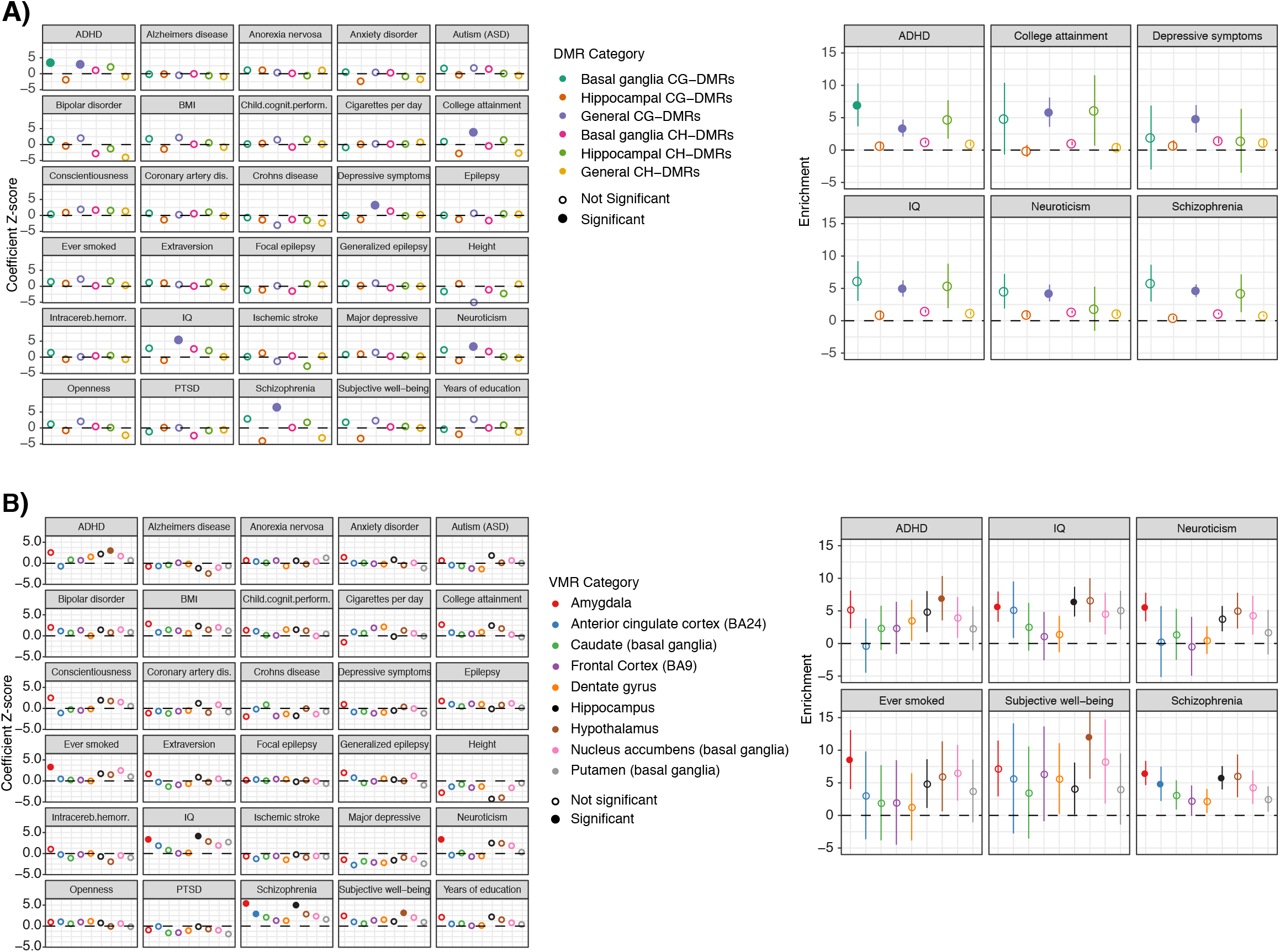
DMRs and VMRs enriched for heritability of neurological traits. Results from running stratified linkage disequilibrium score regression using 30 GWAS traits with the (A) DMRs and (B) VMRs identified in this study and 97 baseline features. Z-scores for each trait are shown (left) and enrichments +/- 2 standard errors are shown (right) only for traits with at least one significant association.

### Supplementary Tables

1. Table S1 Sample phenotype data.
2. Table S2 WGBS sequencing statistics.
3. Table S3 Samples failing genotype quality control check.
4. Table S4 CG-DMRs identified among neuronal samples isolated from all 8 brain regions.
5. Table S5 CG-DMRs identified among the 5 brain region groups (cortical, basal ganglia, hypothalamus, hippocampus, amygdala).
6. Table S6 Large blocks of differential CpG methylation identified among the 5 brain region groups (cortical, basal ganglia, hypothalamus, hippocampus, amygdala).
7. Table S7 CG-DMRs identified among neuronal samples isolated from the basal ganglia tissues (caudate, putamen, nucleus accumbens).
8. Table S8 GREAT analysis of basal ganglia CG-DMRs unique to this analysis and the top 2,000 Hippocampal CG-DMRs.
9. Table S9 CG-DMRs identified among neuronal samples isolated from the two hippocampus tissue groups.
10. Table S10a List of 75 granule cell marker genes (hg38 coordinates). Table S10b Hippocampal CG-DMRs that overlap granule cell marker genes or their promoters (TSS +/- 4 kb).
11. Table S11 CH-DMRs identified among the 5 brain region groups (cortical, basal ganglia, hypothalamus, hippocampus, amygdala).
12. Table S12a CH-DMRs identified among neuronal samples isolated from the basal ganglia tissues. Table S12b CH-DMRs identified among neuronal samples isolated from the two hippocampus tissue groups.
13. Table S13 Lists of VMRs identified in each tissue.

All_VMRs: Union of all VMRs from all tissues (9 brain regions, lung, and thyroid).
Ubiquitious VMRs: VMRs shared across all tissues (9 brain regions, lung, and thyroid). Dentate gyrus all: All VMRs identified in the hippocampus (dentate gyrus) samples.
Dentate gyrus only: Tissue-specific VMRs identified only in the hippocampus (dentate gyrus) samples.
HC only: Tissue-specific VMRs identified only in the hippocampus samples. HC all: All VMRs identified in the hippocampus samples.
AMY only: Tissue-specific VMRs identified only in the amygdala samples. AMY all: All VMRs identified in the amygdala samples.
HYP only: Tissue-specific VMRs identified only in the hypothalamus samples. HYP all: All VMRs identified in the hypothalamus samples.
NAcc only: Tissue-specific VMRs identified only in the nucleus accumbens samples. NAcc all: All VMRs identified in the nucleus accumbens samples.
PUT only: Tissue-specific VMRs identified only in the putamen samples. PUT all: All VMRs identified in the putamen samples.
BA9 only: Tissue-specific VMRs identified in the frontal cortex (BA9) samples. BA9 all: All VMRs identified in the frontal cortex (BA9) samples.
BA24 only: Tissue-specific VMRs identified in the frontal cortex (BA24) samples. BA24 all: All VMRs identified in the frontal cortex (BA24) samples.
CAU all: All VMRs identified in the caudate samples.
CAU only: Tissue-specific VMRs identified only in the caudate samples. Thyroid only: Tissue-specific VMRs identified only in the thyroid samples. Thyroid all: All VMRs identified in the thyroid samples.
Lung only: Tissue-specific VMRs identified only in the lung samples. Lung all: All VMRs identified in the lung samples.
14. Table S14 GREAT analysis of VMRs shared only between amygdala and hypothalamus samples.
15. Table S15 Links and references to summary statistics for 30 traits used in stratified linkage disequilibrium score regression analyses.
16. Table S16 Results from stratified linkage disequilibrium score regression analyses using epigenetic features.
17. Table S17 Overlap between VMRs and SNPs at several minor allele frequencies.

## REFERENCES

1. Lister R, Mukamel EA, Nery JR, Urich M, Puddifoot CA, Johnson ND, Lucero J, Huang Y, Dwork AJ, Schultz MD, et al: Global epigenomic reconfiguration during mammalian brain development. Science 2013, 341:1237905.

2. Zhu J, Adli M, Zou JY, Verstappen G, Coyne M, Zhang X, Durham T, Miri M, Deshpande V, De Jager PL, et al: Genome-wide chromatin state transitions associated with developmental and environmental cues. Cell 2013, 152:642–654.

3. Sun MA, Sun Z, Wu X, Rajaram V, Keimig D, Lim J, Zhu H, Xie H: Mammalian Brain Development is Accompanied by a Dramatic Increase in Bipolar DNA Methylation. Sci Rep 2016, 6:32298.

4. Price AJ, Collado-Torres L, Ivanov NA, Xia W, Burke EE, Shin JH, Tao R, Ma L, Jia Y, Hyde TM, et al: Divergent neuronal DNA methylation patterns across human cortical development reveal critical periods and a unique role of CpH methylation. Genome Biol 2019, 20:196.

5. Pidsley R, Viana J, Hannon E, Spiers H, Troakes C, Al-Saraj S, Mechawar N, Turecki G, Schalkwyk LC, Bray NJ, Mill J: Methylomic profiling of human brain tissue supports a neurodevelopmental origin for schizophrenia. Genome Biol 2014, 15:483.

6. Hannon E, Spiers H, Viana J, Pidsley R, Burrage J, Murphy TM, Troakes C, Turecki G, O’Donovan MC, Schalkwyk LC, et al: Methylation QTLs in the developing brain and their enrichment in schizophrenia risk loci. Nat Neurosci 2016, 19:48–54.

7. Hoffmann A, Sportelli V, Ziller M, Spengler D: Epigenomics of Major Depressive Disorders and Schizophrenia: Early Life Decides. Int J Mol Sci 2017, 18.

8. Jaffe AE, Gao Y, Deep-Soboslay A, Tao R, Hyde TM, Weinberger DR, Kleinman JE: Mapping DNA methylation across development, genotype and schizophrenia in the human frontal cortex. Nat Neurosci 2016, 19:40–47.

9. Karpova NN, Sales AJ, Joca SR: Epigenetic Basis of Neuronal and Synaptic Plasticity. Curr Top Med Chem 2017, 17:771–793.

10. Tognini P, Napoli D, Pizzorusso T: Dynamic DNA methylation in the brain: a new epigenetic mark for experience-dependent plasticity. Front Cell Neurosci 2015, 9:331.

11. Fullard JF, Hauberg ME, Bendl J, Egervari G, Cirnaru MD, Reach SM, Motl J, Ehrlich ME, Hurd YL, Roussos P: An atlas of chromatin accessibility in the adult human brain. Genome Res 2018.

12. Rizzardi LF, Hickey PF, Rodriguez DiBlasi V, Tryggvadottir R, Callahan CM, Idrizi A, Hansen KD, Feinberg AP: Neuronal brain-region-specific DNA methylation and chromatin accessibility are associated with neuropsychiatric trait heritability. Nat Neurosci 2019, 22:307–316.

13. Wang D, Liu S, Warrell J, Won H, Shi X, Navarro FCP, Clarke D, Gu M, Emani P, Yang YT, et al: Comprehensive functional genomic resource and integrative model for the human brain. Science 2018, 362.

14. Li M, Santpere G, Imamura Kawasawa Y, Evgrafov OV, Gulden FO, Pochareddy S, Sunkin SM, Li Z, Shin Y, Zhu Y, et al: Integrative functional genomic analysis of human brain development and neuropsychiatric risks. Science 2018, 362.

15. Kozlenkov A, Li J, Apontes P, Hurd YL, Byne WM, Koonin EV, Wegner M, Mukamel EA, Dracheva S: A unique role for DNA (hydroxy)methylation in epigenetic regulation of human inhibitory neurons. Sci Adv 2018, 4:eaau6190.

16. Aguet F, Barbeira AN, Bonazzola R, Brown A, Castel SE, Jo B, Kasela S, Kim-Hellmuth S, Liang Y, Oliva M, et al: The GTEx Consortium atlas of genetic regulatory effects across human tissues. bioRxiv 2019:787903.

17. Kim-Hellmuth S, Aguet F, Oliva M, Muñoz-Aguirre M, Wucher V, Kasela S, Castel SE, Hamel AR, Viñuela A, Roberts AL, et al: Cell type specific genetic regulation of gene expression across human tissues. bioRxiv 2019:806117.

18. Consortium G: The Genotype-Tissue Expression (GTEx) project. Nat Genet 2013, 45:580–585.

19. Consortium G: Human genomics. The Genotype-Tissue Expression (GTEx) pilot analysis: multitissue gene regulation in humans. Science 2015, 348:648–660.

20. eGTExProject: Enhancing GTEx by bridging the gaps between genotype, gene expression, and disease. Nat Genet 2017, 49:1664–1670.

21. Jaffe AE, Feinberg AP, Irizarry RA, Leek JT: Significance analysis and statistical dissection of variably methylated regions. Biostatistics 2012, 13:166–178.

22. Feinberg AP, Irizarry RA, Fradin D, Aryee MJ, Murakami P, Aspelund T, Eiriksdottir G, Harris TB, Launer L, Gudnason V, Fallin MD: Personalized epigenomic signatures that are stable over time and covary with body mass index. Sci Transl Med 2010, 2:49ra67.

23. Kirsch L, Chechik G: On Expression Patterns and Developmental Origin of Human Brain Regions. PLoS Comput Biol 2016, 12:e1005064.

24. Vermunt MW, Reinink P, Korving J, de Bruijn E, Creyghton PM, Basak O, Geeven G, Toonen PW, Lansu N, Meunier C, et al: Large-scale identification of coregulated enhancer networks in the adult human brain. Cell Rep 2014, 9:767–779.

25. Roadmap Epigenomics C, Kundaje A, Meuleman W, Ernst J, Bilenky M, Yen A, Heravi-Moussavi A, Kheradpour P, Zhang Z, Wang J, et al: Integrative analysis of 111 reference human epigenomes. Nature 2015, 518:317–330.

26. Abad MA, Enguita M, DeGregorio-Rocasolano N, Ferrer I, Trullas R: Neuronal pentraxin 1 contributes to the neuronal damage evoked by amyloid-beta and is overexpressed in dystrophic neurites in Alzheimer’s brain. J Neurosci 2006, 26:12735–12747.

27. Koob GF, Volkow ND: Neurobiology of addiction: a neurocircuitry analysis. Lancet Psychiatry 2016, 3:760–773.

28. McLean CY, Bristor D, Hiller M, Clarke SL, Schaar BT, Lowe CB, Wenger AM, Bejerano G: GREAT improves functional interpretation of cis-regulatory regions. Nat Biotechnol 2010, 28:495–501.

29. Cembrowski MS, Wang L, Sugino K, Shields BC, Spruston N: Hipposeq: a comprehensive RNA-seq database of gene expression in hippocampal principal neurons. Elife 2016, 5:e14997.

30. Mancarci BO, Toker L, Tripathy SJ, Li B, Rocco B, Sibille E, Pavlidis P: Cross-Laboratory Analysis of Brain Cell Type Transcriptomes with Applications to Interpretation of Bulk Tissue Data. eNeuro 2017, 4.

31. Jaffe AE, Hoeppner DJ, Saito T, Blanpain L, Ukaigwe J, Burke EE, Collado-Torres L, Tao R, Tajinda K, Maynard KR, et al: Profiling gene expression in the human dentate gyrus granule cell layer reveals insights into schizophrenia and its genetic risk. Nat Neurosci 2020.

32. Mo A, Mukamel EA, Davis FP, Luo C, Henry GL, Picard S, Urich MA, Nery JR, Sejnowski TJ, Lister R, et al: Epigenomic Signatures of Neuronal Diversity in the Mammalian Brain. Neuron 2015, 86:1369–1384.

33. Li P, Marshall L, Oh G, Jakubowski JL, Groot D, He Y, Wang T, Petronis A, Labrie V: Epigenetic dysregulation of enhancers in neurons is associated with Alzheimer’s disease pathology and cognitive symptoms. Nat Commun 2019, 10:2246.

34. Stroud H, Su SC, Hrvatin S, Greben AW, Renthal W, Boxer LD, Nagy MA, Hochbaum DR, Kinde B, Gabel HW, Greenberg ME: Early-Life Gene Expression in Neurons Modulates Lasting Epigenetic States. Cell 2017, 171:1151–1164 e1116.

35. Keown CL, Berletch JB, Castanon R, Nery JR, Disteche CM, Ecker JR, Mukamel EA: Allele-specific non-CG DNA methylation marks domains of active chromatin in female mouse brain. Proc Natl Acad Sci U S A 2017, 114:E2882–E2890.

36. Kvartsberg H, Lashley T, Murray CE, Brinkmalm G, Cullen NC, Hoglund K, Zetterberg H, Blennow K, Portelius E: The intact postsynaptic protein neurogranin is reduced in brain tissue from patients with familial and sporadic Alzheimer’s disease. Acta Neuropathol 2019, 137:89–102.

37. Ziller MJ, Gu H, Muller F, Donaghey J, Tsai LT, Kohlbacher O, De Jager PL, Rosen ED, Bennett DA, Bernstein BE, et al: Charting a dynamic DNA methylation landscape of the human genome. Nature 2013, 500:477–481.

38. Garg P, Joshi RS, Watson C, Sharp AJ: A survey of inter-individual variation in DNA methylation identifies environmentally responsive co-regulated networks of epigenetic variation in the human genome. PLoS Genet 2018, 14:e1007707.

39. Gunasekara CJ, Scott CA, Laritsky E, Baker MS, MacKay H, Duryea JD, Kessler NJ, Hellenthal G, Wood AC, Hodges KR, et al: A genomic atlas of systemic interindividual epigenetic variation in humans. Genome Biol 2019, 20:105.

40. Rakyan VK, Blewitt ME, Druker R, Preis JI, Whitelaw E: Metastable epialleles in mammals. Trends Genet 2002, 18:348–351.

41. Zheng SC, Breeze CE, Beck S, Dong D, Zhu T, Ma L, Ye W, Zhang G, Teschendorff AE: EpiDISH web server: Epigenetic Dissection of Intra-Sample-Heterogeneity with online GUI. Bioinformatics 2019.

42. Kozlenkov A, Roussos P, Timashpolsky A, Barbu M, Rudchenko S, Bibikova M, Klotzle B, Byne W, Lyddon R, Di Narzo AF, et al: Differences in DNA methylation between human neuronal and glial cells are concentrated in enhancers and non-CpG sites. Nucleic Acids Res 2014, 42:109–127.

43. Etkin A, Prater KE, Schatzberg AF, Menon V, Greicius MD: Disrupted amygdalar subregion functional connectivity and evidence of a compensatory network in generalized anxiety disorder. Arch Gen Psychiatry 2009, 66:1361–1372.

44. Bach DR, Behrens TE, Garrido L, Weiskopf N, Dolan RJ: Deep and superficial amygdala nuclei projections revealed in vivo by probabilistic tractography. J Neurosci 2011, 31:618–623.

45. Saygin ZM, Kliemann D, Iglesias JE, van der Kouwe AJW, Boyd E, Reuter M, Stevens A, Van Leemput K, McKee A, Frosch MP, et al: High-resolution magnetic resonance imaging reveals nuclei of the human amygdala: manual segmentation to automatic atlas. Neuroimage 2017, 155:370–382.

46. Abivardi A, Bach DR: Deconstructing white matter connectivity of human amygdala nuclei with thalamus and cortex subdivisions in vivo. Hum Brain Mapp 2017, 38:3927–3940.

47. Wu YE, Pan L, Zuo Y, Li X, Hong W: Detecting Activated Cell Populations Using Single-Cell RNA-Seq. Neuron 2017, 96:313–329 e316.

48. Aubrey KR: Presynaptic control of inhibitory neurotransmitter content in VIAAT containing synaptic vesicles. Neurochem Int 2016, 98:94–102.

49. Salatino-Oliveira A, Rohde LA, Hutz MH: The dopamine transporter role in psychiatric phenotypes. Am J Med Genet B Neuropsychiatr Genet 2018, 177:211–231.

50. Finucane HK, Bulik-Sullivan B, Gusev A, Trynka G, Reshef Y, Loh PR, Anttila V, Xu H, Zang C, Farh K, et al: Partitioning heritability by functional annotation using genome-wide association summary statistics. Nat Genet 2015, 47:1228–1235.

51. Fatemi SH, Stary JM, Earle JA, Araghi-Niknam M, Eagan E: GABAergic dysfunction in schizophrenia and mood disorders as reflected by decreased levels of glutamic acid decarboxylase 65 and 67 kDa and Reelin proteins in cerebellum. Schizophr Res 2005, 72:109–122.

52. Heckers S, Stone D, Walsh J, Shick J, Koul P, Benes FM: Differential hippocampal expression of glutamic acid decarboxylase 65 and 67 messenger RNA in bipolar disorder and schizophrenia. Arch Gen Psychiatry 2002, 59:521–529.

53. Moyer CE, Delevich KM, Fish KN, Asafu-Adjei JK, Sampson AR, Dorph-Petersen KA, Lewis DA, Sweet RA: Reduced glutamate decarboxylase 65 protein within primary auditory cortex inhibitory boutons in schizophrenia. Biol Psychiatry 2012, 72:734–743.

54. Visel A, Minovitsky S, Dubchak I, Pennacchio LA: VISTA Enhancer Browser--a database of tissue-specific human enhancers. Nucleic Acids Res 2007, 35:D88–92.

55. Schulz H, Ruppert AK, Herms S, Wolf C, Mirza-Schreiber N, Stegle O, Czamara D, Forstner AJ, Sivalingam S, Schoch S, et al: Genome-wide mapping of genetic determinants influencing DNA methylation and gene expression in human hippocampus. Nat Commun 2017, 8:1511.

56. Do C, Lang CF, Lin J, Darbary H, Krupska I, Gaba A, Petukhova L, Vonsattel JP, Gallagher MP, Goland RS, et al: Mechanisms and Disease Associations of Haplotype-Dependent Allele-Specific DNA Methylation. Am J Hum Genet 2016, 98:934–955.

57. Wen L, Li X, Yan L, Tan Y, Li R, Zhao Y, Wang Y, Xie J, Zhang Y, Song C, et al: Whole-genome analysis of 5-hydroxymethylcytosine and 5-methylcytosine at base resolution in the human brain. Genome Biol 2014, 15:R49.

58. Nestor CE, Ottaviano R, Reddington J, Sproul D, Reinhardt D, Dunican D, Katz E, Dixon JM, Harrison DJ, Meehan RR: Tissue type is a major modifier of the 5-hydroxymethylcytosine content of human genes. Genome Res 2012, 22:467–477.

59. Carithers LJ, Ardlie K, Barcus M, Branton PA, Britton A, Buia SA, Compton CC, DeLuca DS, Peter-Demchok J, Gelfand ET, et al: A Novel Approach to High-Quality Postmortem Tissue Procurement: The GTEx Project. Biopreserv Biobank 2015, 13:311–319.

60. Krueger F, Andrews SR: Bismark: a flexible aligner and methylation caller for Bisulfite-Seq applications. Bioinformatics 2011, 27:1571–1572.

61. Hansen KD, Langmead B, Irizarry RA: BSmooth: from whole genome bisulfite sequencing reads to differentially methylated regions. Genome Biol 2012, 13:R83.

62. Moran S, Arribas C, Esteller M: Validation of a DNA methylation microarray for 850,000 CpG sites of the human genome enriched in enhancer sequences. Epigenomics 2016, 8:389–399.

63. Harrow J, Frankish A, Gonzalez JM, Tapanari E, Diekhans M, Kokocinski F, Aken BL, Barrell D, Zadissa A, Searle S, et al: GENCODE: the reference human genome annotation for The ENCODE Project. Genome Res 2012, 22:1760–1774.

64. Kuhn RM, Haussler D, Kent WJ: The UCSC genome browser and associated tools. Brief Bioinform 2013, 14:144–161.

65. Rosenbloom KR, Armstrong J, Barber GP, Casper J, Clawson H, Diekhans M, Dreszer TR, Fujita PA, Guruvadoo L, Haeussler M, et al: The UCSC Genome Browser database: 2015 update. Nucleic Acids Res 2015, 43:D670–681.

66. Durinck S, Moreau Y, Kasprzyk A, Davis S, De Moor B, Brazma A, Huber W: BioMart and Bioconductor: a powerful link between biological databases and microarray data analysis. Bioinformatics 2005, 21:3439–3440.

67. Durinck S, Spellman PT, Birney E, Huber W: Mapping identifiers for the integration of genomic datasets with the R/Bioconductor package biomaRt. Nat Protoc 2009, 4:1184–1191.

68. Lawrence M, Gentleman R, Carey V: rtracklayer: an R package for interfacing with genome browsers. Bioinformatics 2009, 25:1841–1842.

69. Team RC: R: A language and environment for statistical computing. R Foundation for Statistical Computing, Vienna, Austria. 2016. 2016.

70. Chang CC, Chow CC, Tellier LC, Vattikuti S, Purcell SM, Lee JJ: Second-generation PLINK: rising to the challenge of larger and richer datasets. Gigascience 2015, 4:7.

71. Obenchain V, Lawrence M, Carey V, Gogarten S, Shannon P, Morgan M: VariantAnnotation: a Bioconductor package for exploration and annotation of genetic variants. Bioinformatics 2014, 30:2076–2078.

72. Bulik-Sullivan BK, Loh PR, Finucane HK, Ripke S, Yang J, Schizophrenia Working Group of the Psychiatric Genomics C, Patterson N, Daly MJ, Price AL, Neale BM: LD Score regression distinguishes confounding from polygenicity in genome-wide association studies. Nat Genet 2015, 47:291–295.

73. Holm S: A Simple Sequentially Rejective Multiple Test Procedure. Scandnavian Journal of Statistics 1979, 6:65–70.

74. Gentleman RC, Carey VJ, Bates DM, Bolstad B, Dettling M, Dudoit S, Ellis B, Gautier L, Ge Y, Gentry J, et al: Bioconductor: open software development for computational biology and bioinformatics. Genome Biol 2004, 5:R80.

75. Lawrence M, Huber W, Pages H, Aboyoun P, Carlson M, Gentleman R, Morgan MT, Carey VJ: Software for computing and annotating genomic ranges. PLoS Comput Biol 2013, 9:e1003118.

76. Wickham H: ggplot2: elegant graphics for data analysis. Springer; 2016.

